# Fos expression in the periaqueductal gray, but not the ventromedial hypothalamus, is correlated with psychosocial stress-induced cocaine-seeking behavior in rats

**DOI:** 10.1101/2025.01.22.634146

**Authors:** Nicole M. Hinds, Ireneusz D. Wojtas, Desta M. Pulley, Stephany J. McDonald, Colton D. Spencer, Milena Sudarikov, Nicole E. Hubbard, Colin M. Kulick-Soper, Samantha de Guzman, Sara Hayden, Jessica J. Debski, Bianca Patel, Douglas P. Fox, Daniel F. Manvich

## Abstract

Psychosocial stressors are known to promote cocaine craving and relapse in humans but are infrequently employed in preclinical relapse models. Consequently, the underlying neural circuitry by which these stressors drive cocaine seeking has not been thoroughly explored. Using Fos expression analyses, we sought to examine whether the ventromedial hypothalamus (VMH) or periaqueductal gray (PAG), two critical components of the brain’s hypothalamic defense system, are activated during psychosocial stress-induced cocaine seeking. Adult male and female rats self-administered cocaine (0.5 mg/kg/inf IV, fixed-ratio 1 schedule, 2 h/session) over 20 sessions. On sessions 11, 14, 17, and 20, a tactile cue was present in the operant chamber that signaled impending social defeat stress (n=16, 8/sex), footshock stress (n=12, 6/sex), or a no-stress control condition (n=12, 6/sex) immediately after the session’s conclusion. Responding was subsequently extinguished, and rats were tested for reinstatement of cocaine seeking during re-exposure to the tactile cue that signaled their impending stress/no-stress post-session event. All experimental groups displayed significant reinstatement of cocaine seeking, but Fos analyses indicated that neural activity within the rostrolateral PAG (rPAGl) was selectively correlated with cocaine-seeking magnitude in the socially-defeated rats. rPAGl activation was also associated with active-defense coping behaviors during social defeat encounters and with Fos expression in prelimbic prefrontal cortex and orexin-negative cells of the lateral hypothalamus/perifornical area in males, but not females. These findings suggest a potentially novel role for the rPAGl in psychosocial stress-induced cocaine seeking, perhaps in a sex-dependent manner.

## 1. INTRODUCTION

The risk of relapse in individuals with Cocaine Use Disorder (CUD) can persist despite extended periods of abstinence [1,2]. Among the factors that reliably elicits cocaine craving and precipitates relapse in CUD patients is acute exposure to psychosocial stress [3–8]. For example, presentation of experimenter-delivered psychosocial stressors such as the Trier Social Stress Task or personalized scripts/imagery of past socially-stressful experiences can increase self-reported cocaine craving, with stronger levels of stress-induced craving predictive of greater relapse vulnerability [9–12]. Moreover, perceived social/emotional distress in the non-laboratory naturalistic environment has in some studies been reported to precede cocaine craving and relapse episodes [3,13–15]. These findings suggest that exposure to psychosocial stress may serve as a potentially important contributor to cocaine relapse susceptibility [4,6–8].

Stress-induced cocaine relapse is frequently modeled in experimental animals using the reinstatement procedure, in which previously-extinguished cocaine-maintained operant responding (i.e., self-administration) or cocaine-conditioned place preference is re-established following acute exposure to a stressor [16–19]. Studies combining stress-induced reinstatement procedures with neural circuit interrogations have provided key insights into the neurochemicals and neural substrates underlying stress-induced drug-seeking responses [16,17,20–23]. With regard specifically to the stress-induced reinstatement of operant cocaine-seeking behavior, functional roles have been identified for several neuropharmacological effectors, including but not limited to norepinephrine (NE) neurotransmission within the central nucleus of the amygdala (CeA) and bed nucleus of stria terminalis (BNST) [24,25], corticotropin-releasing factor (CRF) receptor activation within the BNST [25–27] and ventral tegmental area (VTA) [25,28–30], dopaminergic and endocannabinoid signaling within the prelimbic prefrontal cortex (plPFC) [30–33], and glutamatergic neurotransmission within the nucleus accumbens (NAc) [32]. However, these findings are derived from studies employing nonsocial stressors to elicit cocaine-seeking responses, typically achieved via systemic or central administration of anxiogenic compounds (e.g., yohimbine, CRF, CRF receptor agonists) or exposure to an inescapable physical stressor (e.g., intermittent footshock). It therefore remains unknown whether the aforementioned mechanisms and substrates play a functional role in psychosocial stress-induced drug-seeking responses or, alternatively, if psychosocial stressors may elicit drug seeking via the engagement of distinct neural circuits. The possibility that psychosocial stressors and nonsocial stressors might elicit drug-seeking behavior through different mechanisms is supported by studies in rodents demonstrating that behavioral responses to nonsocial stressors (e.g., yohimbine, footshock, injury/pain) and psychosocial stressors are largely mediated by non-overlapping neural pathways [34–38]. More specifically, conspecific social defeat stress (SDS) and predator exposure, the two most commonly studied psychosocial stressors in rodents, evoke defensive behaviors and physiological changes via the activation of parallel circuits within a medial hypothalamic defense system comprised of topographically-organized projections from hypothalamic nuclei (ventromedial nucleus, VMH; dorsal premammillary nucleus, PMD) to the midbrain periaqueductal gray (PAG) [34,37–41].

The VMH has well-established roles in several homeostatic and reproductive processes but has importantly been implicated in the generation of innate defensive behaviors as well as emotional and learned responses to social threats [42–45]. There is abundant evidence indicating that defensive reactions to predators and aggressive conspecifics in rodents are dependent upon neural activation of the dorsomedial and ventrolateral aspects of the VMH, respectively (VMHdm, VMHvl) [34,40]. For example, expression of the immediate early gene product Fos, a marker of cellular activation [46–48], is selectively increased in the VMHdm of rodents following exposure to predatory threat but not to an aggressive conspecific, while the opposite is true for the VMHvl [34,49–51]. Similarly, studies employing electrophysiological recordings or calcium imaging have found that neurons in the VMHdm encode defensive reactions to predatory threat, whereas VMHvl neurons increase their firing during conspecific aggressive encounters [50–55]. Electrical or optogenetic stimulation of the VMH evokes defensive behaviors in rodents [42,43,50,51], while lesions or transient inhibition of the VMH attenuates behavioral responses to predatory and conspecific aggressive stimuli in a subregion-specific manner [34,49–51]. Importantly, nonsocial stressors do not appreciably activate the VMH or PMD [34], and inhibition or lesion of these hypothalamic nuclei does not impact behavioral responses to footshock [34,56], indicating that the capacity of the VMH and PMD to engender defensive behavioral repertoires is selective for social stressors.

The VMHdm and VMHvl in turn orchestrate their psychosocial stress-induced defensive responses via direct projections to the PAG as well as via indirect projections through a VMH-PMD-PAG circuit [38,57,58]. The PAG is a highly-conserved midbrain region perhaps best known for its involvement in the descending analgesia system and anxiety/panic processes [59–62], but is also well-established to coordinate the expression of a complex array of behavioral and physiological responses to both social and nonsocial stressors, thus serving as a final common output among various stress-coping systems within the brain [37,40,41,63]. For example, Fos expression in the PAG is increased by predator threat and conspecific social aggression [38,64–67], but also by exposure to footshock [68,69], administration of anxiogenic drugs [70], and in response to several physiological stressors (e.g., hypoxia, hypercapnia) [71,72]. Furthermore, activation of the PAG is notably increased during withdrawal from several drugs of abuse [73–77]. Stimulation of the PAG is sufficient to induce a variety of defensive responses in rodents that can range from passive (e.g., freezing) to active (e.g., escape, aggression) coping behaviors, depending upon the intensity and subregional focus of the stimulation [42,78–84]. This diversity of PAG-mediated behavioral and physiological effects is based upon its segmentation into four longitudinal columns: dorsomedial, dorsolateral, lateral, and ventrolateral [63,82,84,85]. Activation of the dorsal and lateral subregions typically induces sympathetic activation and active defensive behaviors including escape and self-defense/aggression, while stimulation of the ventrolateral subregion is associated with passive coping responses (i.e., freezing) and quiescence [37,82,85,86]. Finally, in contrast to the VMH and PMD, inhibition of PAG activity attenuates defensive behaviors in response to both social stressors [34,45] and nonsocial stressors [79,87–92].

The functional roles of the VMH and PAG in producing adaptive responses to danger appears to be well-conserved in nonhuman primates and humans. In the common marmoset, predator threat induces neuronal activation in the VMHdm [39], while electrical stimulation of the VMH elicits an intense defensive state [93]. Similarly in humans, predator approach in a virtual reality environment is associated with increased activation of the PAG [94], and electrical stimulation of the VMH, dPAG, or lPAG evokes feelings of doom and terror, sensations of being chased, and autonomic activation that are highly reminiscent of the symptomology of panic attacks [95–97]. Yet despite this evidence that the VMH and PAG underlie stress-evoked defensive behaviors across species, their potential involvement in the expression of stress-induced drug-seeking behaviors has received little attention to date. Some support for this possibility comes from human neuroimaging studies suggesting that hyperactivation of the PAG (through the proposed weakening of top-down inhibition from frontal cortical regions) is associated with greater severity of alcohol use, cocaine use, and cocaine craving [98–100]. These clinical reports align with findings in mice that optogenetic inhibition of medial prefrontal cortical input to the dPAG drives compulsive alcohol drinking, whereas optogenetic activation of this projection is sufficient to suppress alcohol drinking [101]. However, no studies to date have explored whether psychosocial stressors may elicit drug seeking through activation of the medial hypothalamic defensive system or its outputs to the dorsal and lateral PAG subregions.

We previously developed a novel model of psychosocial stress-induced reinstatement whereby cocaine-seeking behavior is reinstated in male rats via exposure to a cue signaling impending conspecific SDS [102]. We selected SDS as a psychosocial stressor because it has been suggested to more closely recapitulate the types of stressful psychosocial experiences that may elicit cocaine craving and relapse in human cocaine users [4,103–105]. In addition to demonstrating cocaine seeking responses to SDS-predictive cues, we also found that animals’ predilection towards displaying active coping responses during SDS was associated with greater levels of stress-induced cocaine-seeking magnitude assessed at a later time point [102]. However, the neural circuits mediating the observed psychosocial stress-induced cocaine-seeking response or the association between cocaine-seeking magnitude and prior active-coping action selection were not explored. The primary objectives of the present study were therefore to examine whether neural activation within components of the medial hypothalamic defense system is associated with psychosocial stress-induced cocaine seeking in male and female rats, and to determine whether neural activity in any of these regions during cocaine seeking was correlated with individual propensities to engage in active coping strategies during social conflicts. We focused our investigation on the VMH and the PAG.

## 2. MATERIALS AND METHODS

### 2.1 Animals

Experimental subjects destined for self-administration and reinstatement studies were adult male (200-225 g upon arrival, n=24) and female (175-200 g upon arrival, n=24) Long-Evans rats acquired from Charles River Laboratories Inc. (Wilmington, MA) or Envigo (Indianapolis, IN). The animals were housed in polycarbonate cages (43 x 24 x 20 cm) under a 12-h reverse light-dark cycle (lights off at 9:00am) with *ad libitum* access to food and water and provision of enrichment. Experimental subjects were initially pair-housed or triple-housed upon arrival in same-sex pairs or triplets and were allowed to acclimate to the facility for one week prior to onset of experiments. A separate cohort of adult male (225-250 g upon arrival) and female (175-200 g upon arrival) Long-Evans rats served as resident-aggressors in social defeat stress studies. These animals were initially housed in same-sex pairs or triplets in polycarbonate cages as described above. Male resident-aggressors were screened for appropriate levels of aggression prior to use in experiments by temporarily removing their cagemate and introducing a novel adult male Sprague-Dawley rat (175-200 g upon arrival, Charles River Laboratories or Envigo) into their home cage. The animals were then allowed to interact for 4-10 min. A trained experimenter carefully observed the behavior exhibited by the potential resident-aggressor, and animals that displayed inappropriate levels of aggression were eliminated as potential resident-aggressors. Screenings took place on 2-4 occasions separated by at least 2 days. Once male resident-aggressor subjects were identified, they were pair-housed with a sexually-receptive female rat in a larger polycarbonate enclosure (68 x 56 x 39 cm) for the remainder of the study. All procedures were conducted in accordance with the NIH Guide for the Care and Use of Laboratory Animals and were approved by the Rowan University Institutional Animal Care and Use Committee. Behavioral studies took place between 09:30am and 05:00pm.

### 2.2 Food training

Following acclimation, rats destined for cocaine self-administration were first trained to lever-press for 45mg food pellets (Bio-Serv, Flemington, NJ) in operant chambers housed within ventilated, sound-attenuating cubicles (Med Associates Inc,. St. Albans, VT). Training sessions (maximum 4 h duration) began with the extension of two levers (left, right), one of which was designated as the active lever and the other the inactive lever in a randomized and counterbalanced manner across subjects. A single response on the active lever resulted in a single food pellet being dispensed into a receptacle located equidistant between the two levers. Animals were considered trained to lever-press once they had earned ≥ 75 reinforcers with ≥ 70% selectivity for the active lever during a single session (typically occurring within 1-3 sessions). Once they were deemed food-trained, animals progressed to surgical catheter implantation as described below. If an animal failed to acquire food training within 4 sessions, food training was halted but the animal still underwent catheter implantation and cocaine self-administration because in our experience food-training expedites, but is not required for, subsequent successful acquisition of IV cocaine self-administration.

### 2.3 Surgery

Following food training, animals were surgically prepared with a chronic indwelling intravenous (IV) catheter as described previously [102,106]. Briefly, under inhaled isoflurane anesthesia, a custom-fabricated silastic catheter was introduced into the right jugular vein and secured to the vessel using nonabsorbable sutures. The catheter was passed subcutaneously to an infusion port positioned in the mid-scapular region from which an infusion cannula protruded through the skin. Animals received meloxicam (1.0 mg/kg SC) prior to the surgery and either chewable tablets of carprofen (2 mg PO) or injectable meloxicam (1.0 mg/kg SC) post-operatively for 2 days. Beginning on the day of surgery and continuing 5-6 days per week thereafter, catheters were flushed with 0.05 ml gentamicin (4 mg/ml) and locked with 0.1 ml heparin (300 units/ml). When not in use, catheters were protected with a silastic obturator and a stainless-steel dust cap. Beginning on the day of surgery, animals were singly-housed for the remainder of the study and given a minimum of one week recovery prior to the onset of self-administration experiments.

### 2.4 Cocaine self-administration and extinction

Cocaine self-administration was conducted 5-7 days/week in 2-h sessions on a fixed-ratio 1 (FR1) schedule of reinforcement. Each session began with extension of both levers and illumination of a house light located on the opposite wall which served as a discriminative stimulus to signal reinforcer availability. An automated syringe pump (PHM100; Med Associates Inc.) located outside the cubicle that housed the operant chamber was connected via Tygon tubing to a 22-g liquid swivel (Instech, Plymouth Meeting, PA) that was secured to a counterweighted arm over the chamber. The tubing then passed through a metal tether that connected to the vascular access port of the animal. Med-PC V Software (Med Associates) interfaced with each operant chamber to control outputs and record lever presses. Animal weights were entered into the self-administration program prior to each session to calculate pump durations for accuracy of cocaine unit doses.

During self-administration sessions, an active-lever response resulted in an infusion of cocaine (0.5 mg/kg/inf) that was paired with illumination of a cue light above the active lever and termination of the house light for 20-s. During this timeout, active-lever presses were recorded but had no scheduled consequences. Inactive-lever presses were recorded throughout the session but also had no consequences. Daily sessions were terminated if 2 h elapsed or 60 cocaine infusions were earned, whichever occurred first. If the session ended before the 2-h time limit, levers were retracted and cue/house lights were turned off, but the animal remained in the chamber for the remainder of the 2-h session. Rats that failed to acquire cocaine self-administration (n=2 males) or suffered a loss of catheter patency prior to the final cocaine self-administration session (n=1 male, n=2 females) were removed from the study.

Rats self-administered cocaine over 20 sessions as described above and were promptly returned to their home cage immediately after each self-administration session’s conclusion except for sessions 11, 14, 17, and 20. During sessions 11, 14, 17, and 20, self-administration took place within a perforated polycarbonate enclosure (26 x 23 x 18 cm) that was situated within the operant chamber. The enclosure provided a change to the tactile sensation of the self-administration context, from a steel-rod floor and solid walls to a perforated (0.375-inch diameter holes) polycarbonate texture on both the walls and the floor. The enclosure allowed for unobstructed access to levers, view of all stimulus lights, and free movement of the tether and the animal. At the end of these self-administration sessions, rats were removed from the operant chamber and immediately exposed to one of three assigned post-session events: social defeat stress (SDS), footshock stress (FS) or an empty cage no-stress control (EC) as described below. For each animal, their assigned post-session event was the same across sessions 11, 14, 17, and 20, thus the enclosure was conditioned to signal impending exposure only to social defeat stress, footshock stress, or a no-stress control condition. No animals were exposed to more than one type of post-session event.

Beginning on session 21, responding was extinguished in 2-h sessions (5-7 days/week) during which responses on the active or inactive lever were recorded but had no scheduled consequences. Responding was considered extinguished when animals emitted ≤ 15 active-lever responses across three of four consecutive sessions. Animals that failed to satisfy extinction criteria within 40 extinction sessions were removed from the study (n=1 male, n=2 females).

### 2.5 Reinstatement

For each animal, reinstatement testing occurred on the day immediately after extinction criteria were met. During the reinstatement session, rats were re-exposed within the operant chamber to the perforated polycarbonate enclosure that had previously signaled that their assigned post-session event was impending, and responding was measured under extinction conditions for 2 h. A complete timeline for cocaine self-administration, extinction, and reinstatement testing is shown in **Fig. 1**.

**Figure 1:**
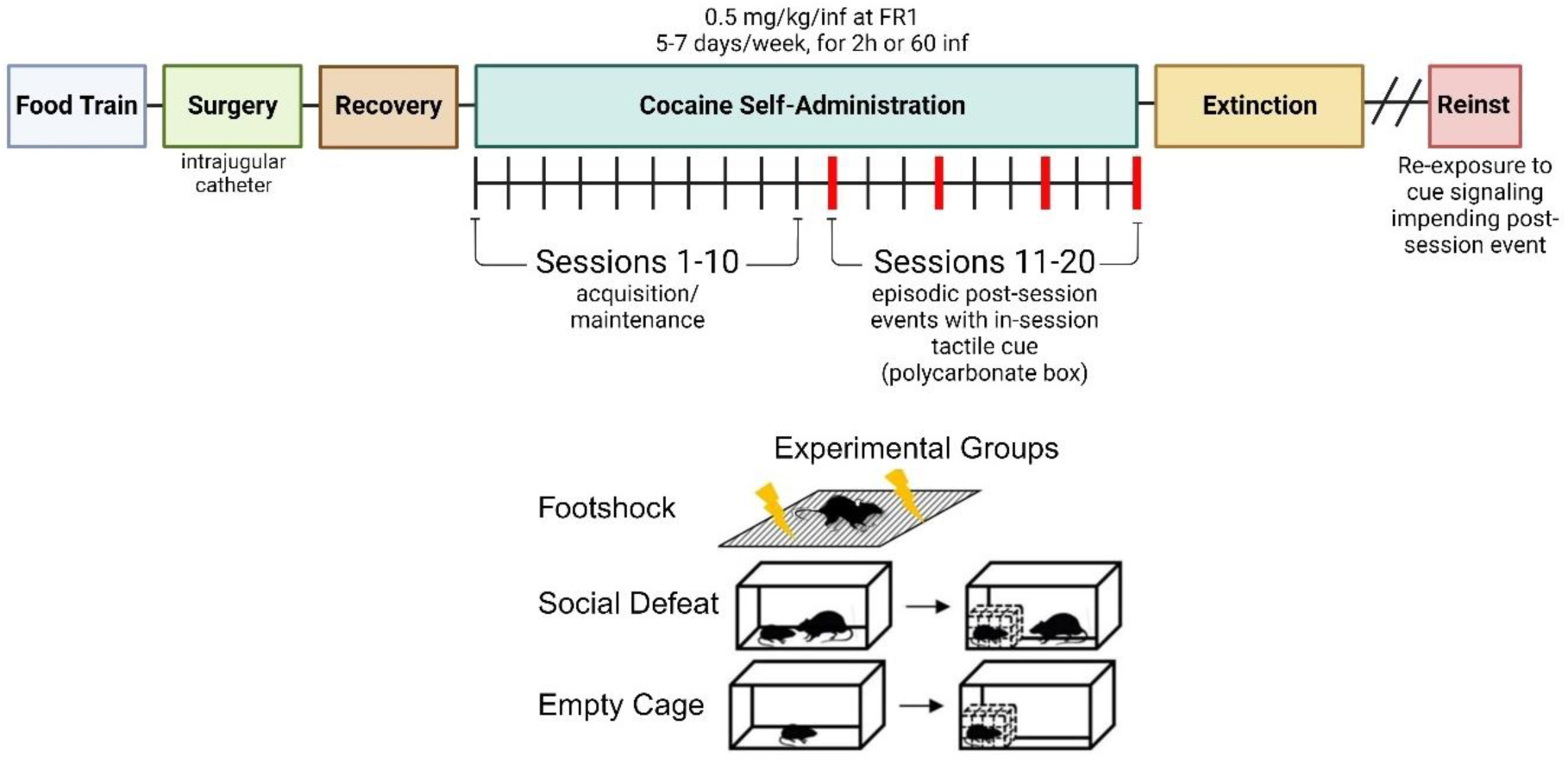
Experimental timeline. Animals self-administered 0.5 mg/kg/inf IV cocaine for 20 sessions under a FR1 schedule of reinforcement. Self-administration during sessions 11, 14, 17, and 20 (designated by red ticks on the timeline) occurred inside a perforated polycarbonate enclosure within the operant chamber and were followed immediately by exposure to social defeat stress (SDS, n=16, 8M/8F), footshock (FS, n=12, 6M/6F), or an empty cage control condition (EC, n=12, 6M/6F). From session 21 onwards, responding in 2-h extinction-training sessions had no scheduled consequences. Reinstatement testing occurred the day after extinction criteria were met. During the reinstatement test, animals were re-exposed to the polycarbonate enclosure that signaled their assigned impending post-session stimulus (SDS, FS, or EC) and were allowed to respond under extinction conditions for 2 h. “Reinst”, reinstatement.

### 2.6 Post-session events

#### 2.6.1 Social defeat stress (SDS)

Immediately following self-administration sessions 11, 14, 17, and 20, rats in the SDS group underwent social defeat stress via the resident-intruder paradigm as described previously [102]. Briefly, “intruder” rats (i.e., cocaine self-administration subjects) were removed from the operant chamber, their catheter was flushed and locked, and they were transported to a housing room adjacent to the operant-behavioral testing room where they were promptly placed into the home cage of a same-sex, conspecific resident-aggressor. When the encounter was a male-male interaction, the female resident-aggressor and any pups were removed from the resident-aggressor’s home cage 2-3 min prior to the intruder’s introduction. When the encounter was a female-female interaction, the male resident-aggressor was removed from the resident-aggressor’s home cage 7-10 min prior to the intruder’s introduction but pups were allowed to remain. Every attempt was made to ensure that female resident-aggressors used to impose conspecific aggression were a dam to pups between PND 3-14 at the time of testing, however in some rare cases pups were slightly older. Direct contact between the intruder and resident-aggressor was stopped if 1) the intruder was bitten twice, 2) the intruder displayed supine submissive posture for 4 consecutive seconds, or 3) 4 minutes elapsed, whichever occurred first. Immediately upon termination of the physical interaction, the intruder was placed back into the same perforated polycarbonate enclosure in which the preceding self-administration session had taken place, and the enclosure was then positioned inside the resident’s home cage for 5 min. This period of protected contact allowed for additional visual, olfactory, and auditory threat from the resident-aggressor without further physical contact. Whenever possible, intruders were exposed to a different resident-aggressor for each of their 4 social defeat stress episodes, however there were rare occurrences when an experimental subject had to be exposed to one resident-aggressor on two occasions.

All social defeat stress episodes were video-recorded and scored using The Observer XT software (Noldus Information Technology, Leesburg, VA) by a trained observer who was blinded to intruder and resident-aggressor identifiers. Scorers quantified the duration of time spent engaged in several defensive behaviors including boxing, push, dominant posture, escape/flight, upright defense, freezing, and supine submissive posture (see *Supplementary Material* for operational definitions of each behavior). In each social defeat stress episode, the percentage of total interactive time engaged in each behavior was calculated, and from these data the following categorical coping scores were generated per episode:

1. “active-defense” coping: sum of % time engaged in boxing, push, and dominant posture
2. “active-avoidance” coping: % time engaged in escape/flight
3. “passive” coping: sum of % time engaged in upright defense, freezing, and supine submissive posture

For each subject, the % time engaged in each individual behavior and each coping category was averaged across the 4 social defeat episodes to generate a final value per animal that was included in statistical analyses.

#### 2.6.2 Footshock (FS)

Immediately following self-administration sessions 11, 14, 17, and 20, rats in the FS group were removed from the operant chamber, their catheter was flushed and locked, and they were then returned to their operant chamber with the polycarbonate enclosure removed, levers retracted, and all lights off. They were then exposed to 15 min of intermittent, unpredictable footshock (20 shocks, 0.5 mA, 0.5 s in duration, variable presentation interval of 3-80 s [24,25,110]) delivered to the electrified floor of the operant chamber. These footshock parameters were chosen on the basis that they reliably reinstate cocaine-seeking behavior in rats [111–113]. Animals were returned to their home cage at the conclusion of the 15-min exposure period.

#### 2.6.3 Empty cage control (EC)

Immediately following self-administration sessions 11, 14, 17, and 20, rats in the EC group were subjected to the same post-session procedure as the rats in the SDS group, with the critical exception that rather than be exposed to a same-sex conspecific aggressive rat, they were instead placed into an uninhabited clean resident-aggressor home cage prepared with fresh bedding. The objective was to approximate all aspects of the SDS condition absent any social threat. EC rats were first allowed to freely explore the clean cage for 4 min. They were then placed back into the same perforated polycarbonate enclosure in which the preceding self-administration session had taken place, and the enclosure was situated within the empty resident cage for a further 5 min. They were then removed from the enclosure and returned to their own home cage.

### 2.7 Tissue preparation

Immediately after the conclusion of the 2-hr reinstatement test, rats were deeply anesthetized via administration of either 10.0 mg/kg ketamine IV delivered through the catheter preparation or, if the catheter was no longer patent, IP administration of a ketamine/xylazine cocktail (100 mg/kg ketamine : 10 mg/kg xylazine), and then expeditiously transported out of the operant-behavioral testing room to a separate laboratory. Under supplementation with 3-5% inhaled isoflurane, rats were transcardially perfused with 150 ml ice-cold 0.1M phosphate-buffered saline (PBS) followed by 250 ml ice-cold 4% paraformaldehyde (PFA) in 0.1M PBS. Brains were removed and postfixed in 4% PFA overnight before performing two sequential submersions in 30% sucrose in 0.1M PBS at 4°C. Brains were then frozen via submersion in dry ice-chilled isopentane and stored at -80°C prior to sectioning. Serial 30 μm coronal sections were collected at every other slice between +4.20 to -10.20 mm anteroposterior (A/P) from bregma and every fourth slice between -10.32 to -15.00 mm A/P from bregma. Sections were stored in cryoprotectant (30% sucrose, 30% ethylene glycol in 0.1M PBS, pH 7.4) at -20°C until immunohistochemical analysis.

### 2.8 Immunohistochemistry

Tissue containing brain regions of interest were removed from cryoprotectant and washed 3 x 10 min in 4 ml of 0.1M PBS. Sections were then blocked for 90 min in 3 ml of buffer containing 5% normal donkey serum (Millipore-Sigma, Burlington, MA), 1% bovine serum albumin (Proliant Biologicals, Ankeny IA) and 94% PBS-T (PBS containing 0.1% Triton® X-100 (VWR Inc., West Chester, PA)) at room temperature. All sections were then incubated in blocking buffer with rabbit anti-Fos primary antibody (1:5000, ab190289, Abcam, Waltham, MA) overnight at room temperature. Sections containing the ventral tegmental area (VTA) were co-incubated with chicken anti-tyrosine hydroxylase (1:1000, ab121013, Abcam), while sections containing the lateral hypothalamus and perifornical area (LH/PfA) were co-incubated with mouse anti-orexin-A (1:1000, sc-80263, Santa Cruz Biotechnology, Dallas, TX). Sections were then washed 3 x 10 min in 4 ml of 0.1M PBS and incubated for 2 h in blocking buffer containing donkey anti-rabbit AlexaFluor™ 488 secondary antibody (1:500, A21206, Invitrogen, Waltham, MA). VTA-containing sections were co-incubated with goat anti-chicken AlexaFluor™ 594 (1:500, A11042, Invitrogen), while LH/PfA containing sections were co-incubated with donkey anti-mouse AlexaFluor™ 647 (1:500, A31571, Invitrogen). Sections lacking VTA or LH/PfA were co-incubated with a fluorescent Nissl stain (1:500, NeuroTrace™ 640/660, Invitrogen) during the secondary antibody incubation period. Sections were then washed 3 x 10 min in 0.1M PBS and subsequently mounted on chromium-dipped microscope slides, air dried, and coverslipped using Vectashield® with DAPI mounting medium (Vector Laboratories, Newark, CA). Slides were stored in a light-protective case at 4°C prior to imaging.

### 2.9 Image acquisition and analysis

Images of brain sections were photographed at 20x magnification using Keyence BZ-X710 and BZ-X810 fluorescence microscopes (Keyence, Itasca, IL). Stacks of 20x image were acquired in each visual field at a 1.0-1.4 μm pitch through the full focal plane of the tissue. Stacked images were subsequently compressed into single two-dimensional images and stitched together using Keyence Analyzer software to create a macro-scale mosaic image of each tissue section. Images were adjusted for brightness and contrast using Adobe Photoshop. Brain regions of interest were outlined based on anatomical landmarks according to the rat brain atlas of Paxinos and Watson [114] and included the dorsomedial and ventrolateral aspects of the ventromedial hypothalamus (VMHdm, VMHvl), dorsomedial, dorsolateral, lateral, and ventrolateral aspects of the rostral/intermediate periaqueductal gray (rPAGdm, rPAGdl, rPAGl, rPAGvl), ventral tegmental area (VTA), lateral hypothalamus and perifornical area (LH/PfA), central amygdala (CeA), and prelimbic (pl) and infralimbic (il) prefrontal cortex (PFC). In each mosaic image, numbers of Fos-expressing cells were manually counted by a trained experimenter blinded to experimental conditions using ImageJ (National Institute of Health) and Adobe Illustrator (Adobe, San Jose, CA) and quantified as the number of Fos-positive cells divided by the area of the brain region, yielding a final value of cells/mm^2^. In most cases, Fos expression within each brain region of interest was counted bilaterally in 2-4 sections per subject. Values were first averaged between hemispheres in each section, and these values were then averaged across sections to generate a single Fos expression value per brain region per subject. On exceptionally rare occasions (< 1% of all data points collected), Fos expression was calculated from a single tissue section or data could not be acquired due to complications with tissue integrity or accidental tissue loss.

### 2.10 Drugs

Cocaine hydrochloride was provided by the National Institute on Drug Abuse Drug Supply Program (RTI International, Research Triangle Park, NC) and dissolved at a concentration of 3.0 mg/ml in bacteriostatic saline. Cocaine solutions were passed through a 0.22 μm filter (Corning Life Sciences, Tewksbury, MA) and stored in sterile vials at 4°C when not in use. All doses are reported as the salt weight.

### 2.11 Statistical analyses

#### Cocaine self-administration

Within each experimental group, the number of active-lever responses, inactive-lever responses, and cocaine infusions earned were first analyzed using two-way mixed-factors ANOVA with sex as the between-subjects variable and session as the within-subjects variable. Subsequent two-way mixed-factors ANOVAs compared each of these measures with experimental group as the between-subjects variable and session as the within-subjects variable, with male and female subjects collapsed together within each group. One subject’s data (n=1 SDS female) was excluded from analyses of responses and earned infusions due to mechanical malfunction of a syringe pump during one self-administration session. Another subject’s data (n=1 EC male) was excluded from the analysis of inactive-lever presses due to observations that the animal was registering high numbers of inactive-lever responses in a non-goal-directed manner (i.e., the subject was repeatedly “bumping” into the inactive lever). Cumulative cocaine intake was compared between groups using a one-way ANOVA.

#### Social defeat stress coping measures in SDS rats

Sex differences in the duration of social defeat stress episodes was analyzed via unpaired t-test. Sex differences in individual coping behaviors or categorial coping scores (“active-defense”, “active-avoidance”, “passive”) were analyzed using Bonferroni-corrected families of unpaired t-tests.

#### Extinction and reinstatement

The rate to satisfy extinction criteria was compared between sexes or experimental groups using an unpaired t-test or one-way ANOVA, respectively. Active-lever and inactive-lever presses emitted during reinstatement tests were first compared between males and females within each experimental group using unpaired t-tests, and then subsequently assessed between experimental groups with male and female subjects collapsed together using a two-way mixed-factors ANOVA with phase (extinction, reinstatement) as the within-subjects variable and experimental group as the between-subjects variable.

#### Fos expression

For each experimental group, Fos expression within a given subcomponent of the medial hypothalamic defense network (VMHdm, VMHvl, rPAGdm, rPAGdl, rPAGl, rPAGvl) or in rPAG-projecting regions of interest (VTA, CeA, LH/PfA, plPFC, ilPFC) was first compared between males and females via Bonferroni-corrected unpaired t-tests. Differences in Fos expression between experimental groups were analyzed via one-way ANOVA. A factor analysis (principal components analysis, PCA) was applied to each experimental group to compare associations between cocaine-seeking behavior (i.e. number of active-lever presses emitted during the reinstatement test) and Fos expression in VMH or rPAG subregions. Because we had previously found associations between social defeat stress-coping behaviors and cocaine-seeking behavior [102], behavioral coping scores from social defeat episodes were also included in the factor analysis for the SDS group. Only factors with an eigenvalue > 1 were extracted. Based on the PCA results, Pearson correlation analyses were used to assess specific associations between cocaine-seeking behavior and Fos expression in rPAG subregions. A separate set of Pearson correlation analyses was performed in each experimental group to determine whether Fos expression in any rPAG subregion was correlated with Fos expression in brain regions with known afferent projections to the rPAG (VTA, CeA, LH/PfA, plPFC, ilPFC). The SDS group was further subdivided by sex, and a Pearon’s correlation network analysis was used to examine relationships among behavioral variables (cocaine-seeking, “active-defense” coping) and Fos expression in rPAGl and upstream sources of afferent input.

When appropriate, significant main effects or interactions in ANOVAs were followed by *post hoc* Holm-Sidak’s pairwise comparisons as specified in the text. ANOVAs, t-tests, and correlation analyses were performed and figures were plotted using GraphPad Prism v10.3 (GraphPad Software Inc., La Jolla, CA). PCA was performed using SPSS Statistics v29.0 (IBM Corp., Armonk, NY). For all statistical analyses, significance was set at *p* < 0.05. Group data are expressed as mean values ± SEM.

## 3. RESULTS

### 3.1 Cocaine self-administration

Cocaine-maintained responding stabilized within the first 3-5 self-administration sessions, with animals responding predominantly on the active lever thereafter (**Fig. 2A**). Two-way ANOVAs comparing males and females within each experimental group (EC, FS, SDS) were performed on measures of active-lever responses, inactive-lever responses, and number of cocaine infusions earned across the 20 self-administration sessions to determine whether biological sex impacted cocaine self-administration under the present conditions. No significant main effect of sex nor a session × sex interaction was detected (*p* > 0.05, data not shown), therefore male and female subjects were collapsed together within each experimental group for subsequent analyses.

**Figure 2:**
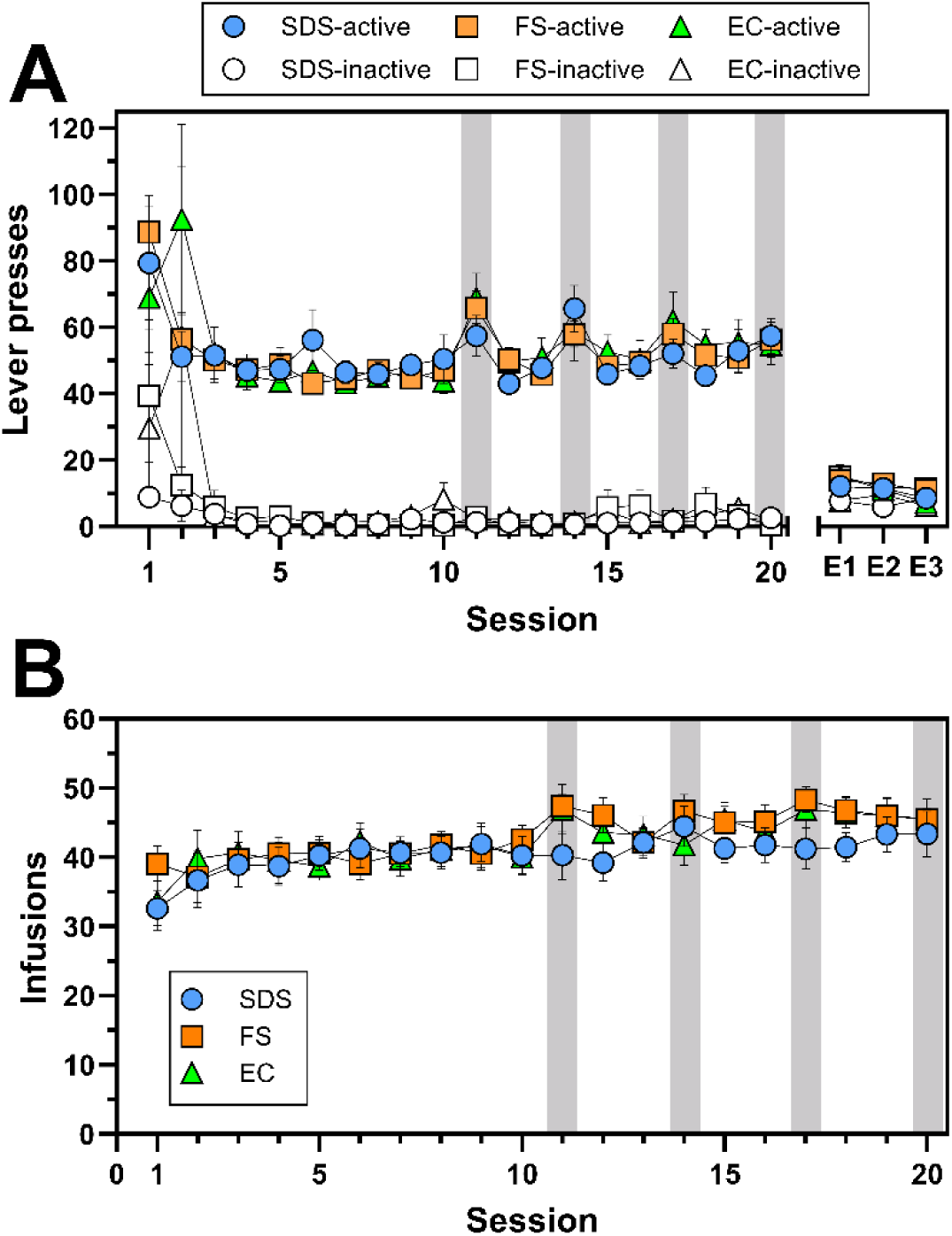
Cocaine self-administration in SDS, FS, and EC rats. **(A)** Active and inactive lever presses (filled and empty symbols, respectively) during cocaine self-administration (sessions 1-20) and the final 3 extinction sessions prior to reinstatement testing (E1-E3) in SDS (n=16, 8M/8F), FS (n=12, 6M/6F), and EC (n=12, 6M/6F) rats. **(B)** Number of cocaine infusions earned across sessions 1-20. Sessions in which the tactile cue and subsequent stress or no-stress post-session events occurred are shaded in gray. Data are presented as mean ± SEM values.

Two-way ANOVA of active-lever responses between sessions 1-20 revealed a main effect of session (*F*_19,684_ = 6.34, *p* < 0.0001) but not group (*F*_2,36_ = 0.10, *p* = 0.91) nor a session × group interaction (*F*_38,684_ = 1.14, *p* = 0.26) (**Fig. 2A**). Two-way ANOVA of inactive-lever responding also indicated a main effect of session (*F*_19,665_ = 3.53, *p* < 0.0001) but not group (*F*_2,35_ = 1.31, *p* = 0.28) nor a session × group interaction (*F*_38,665_ = 1.08, *p* = 0.35) (**Fig. 2A**). Similarly, two-way ANOVA of reinforcers indicated that the number of earned cocaine infusions varied significantly across the 20 self-administration sessions (*F*_19,684_ = 6.21, *p* < 0.0001) but did not differ between groups (*F*_2,36_ = 0.46, *p* = 0.64) nor was there a session × group interaction (*F*_38,684_ = 0.87, *p* = 0.70) (**Fig. 2B**). Accordingly, cumulative cocaine intake did not differ between groups (one-way ANOVA, *F*_2,37_ = 0.12, *p* = 0.89) (**Supplementary Fig. 1**). Collectively, these results indicate that there were no significant differences between the three experimental groups in either cocaine-maintained responding or cocaine intake across the 20 cocaine self-administration sessions.

### 3.2 Behavioral coping responses during social defeat stress episodes (SDS group only)

Rats in the SDS group underwent social defeat stress via the resident-intruder procedure immediately after self-administration sessions 11, 14, 17, and 20. Physical contact between the intruder and the resident-aggressor was allowed to continue for a maximum of 4 min but was terminated earlier if the intruder was pinned in a submissive supine posture for 4 consecutive seconds or if they were bitten twice. The mean duration of social defeat episodes did not differ between sexes (males, 159.7 ± 18.05 s; females, 188.5 ± 14.30 s; *t*_14_ = 1.25, *p* = 0.23). The percentage of time spent engaged in various defensive behaviors during social defeat episodes is shown in **Fig. 3**. On average, animals spent > 75% of the episode engaged in passive behaviors (freezing, upright defense, supine submissive posture). Bonferroni-corrected t-tests revealed that females spent significantly more time engaged in “active-avoidance” coping (i.e., escape/flight behavior) as compared to males (p < 0.05). There were no sex differences detected for any other behaviors.

**Figure 3:**
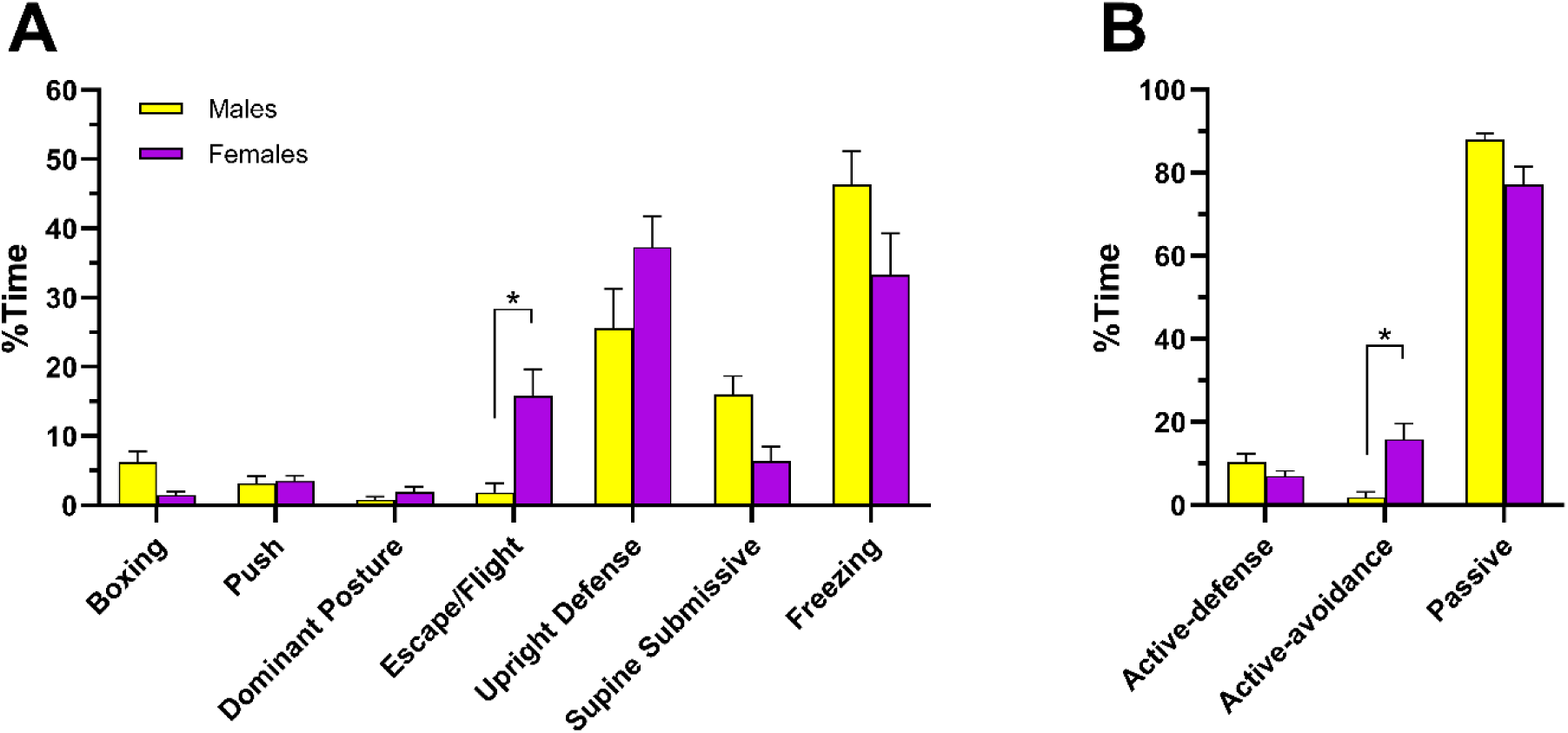
Frequency of behaviors exhibited by SDS rats during social defeat stress episodes. SDS rats (n=16, 8M/8F) were exposed to an aggressive same-sex conspecific immediately after cocaine self-administration sessions 11, 14, 17, and 20. Episodes were video-recorded and the amount of time spent engaged in defensive behaviors were measured. **(A)** Percentages of time male and female rats spent engaged in various defensive behaviors during physical interaction with a resident-aggressor (prior to protected contact). **(B)** Behavioral coping scores derived from categorical sorting and summation of %time values presented in **A**. Data are presented as mean ± SEM values. * *p* < 0.05, males vs. females.

### 3.3 Extinction training

Rats satisfied extinction criteria within 15.83 ± 1.65 (range 5-40) extinction sessions. There were no differences in the rate to reach extinction between the three experimental groups (one-way ANOVA, *F*_2,37_ = 0.27, *p* = 0.77; **Supplementary Fig. 2A**) nor was there a sex difference when the experimental groups were collapsed (*t*_36_ = 0.98, p = 0.34; **Supplementary Fig. 2B**).

### 3.4 Reinstatement

We first examined whether there were sex differences in reinstatement magnitude within any of the three experimental groups (EC, FS, SDS). Independent t-tests comparing males and females on measures of active-lever responding or inactive-lever responding during reinstatement test sessions revealed no significant differences (*p* > 0.05, data not shown). Consequently, males and females were combined within each experimental group for further analyses of reinstatement data.

Levels of reinstated cocaine-seeking behavior produced by re-exposure to the tactile cue (i.e., perforated polycarbonate enclosure) associated with impending SDS, FS, or EC stimulus presentation are shown in **Fig. 4**. Two-way ANOVA comparing active-lever responses between extinction and reinstatement phases revealed a main effect of phase (*F*_1,37_ = 54.59, *p* < 0.0001) but not group (*F*_2,37_ = 2.28, *p* = 0.12) nor was there a phase × group interaction (*F*_2,37_ = 2.06, *p* = 0.14), indicating that re-exposure to the perforated polycarbonate enclosure resulted in significant reinstatement of cocaine-seeking behavior that was equivalent in magnitude between groups (**Fig. 4A**). Notably, levels of active-lever responding during the reinstatement test were comparable to those emitted during prior cocaine self-administration sessions (i.e., during maintenance). Two-way ANOVA of inactive-lever presses between extinction and reinstatement indicated no effect of phase (*F*_1,37_ = 0.24, *p* = 0.63) or group (*F*_2,37_ = 1.64, *p* = 0.21), nor a phase × group interaction (*F*_2,37_ = 0.96, *p* = 0.39) (**Fig. 4B**), indicating that cocaine-seeking behavior during reinstatement test sessions was selectively allocated to the previously-reinforced active lever. For each group, the majority of active-lever responses during reinstatement tests (∼65-70%) were emitted within the first 30 min of the 2-h test session (**Supplementary Fig. 3**).

**Figure 4:**
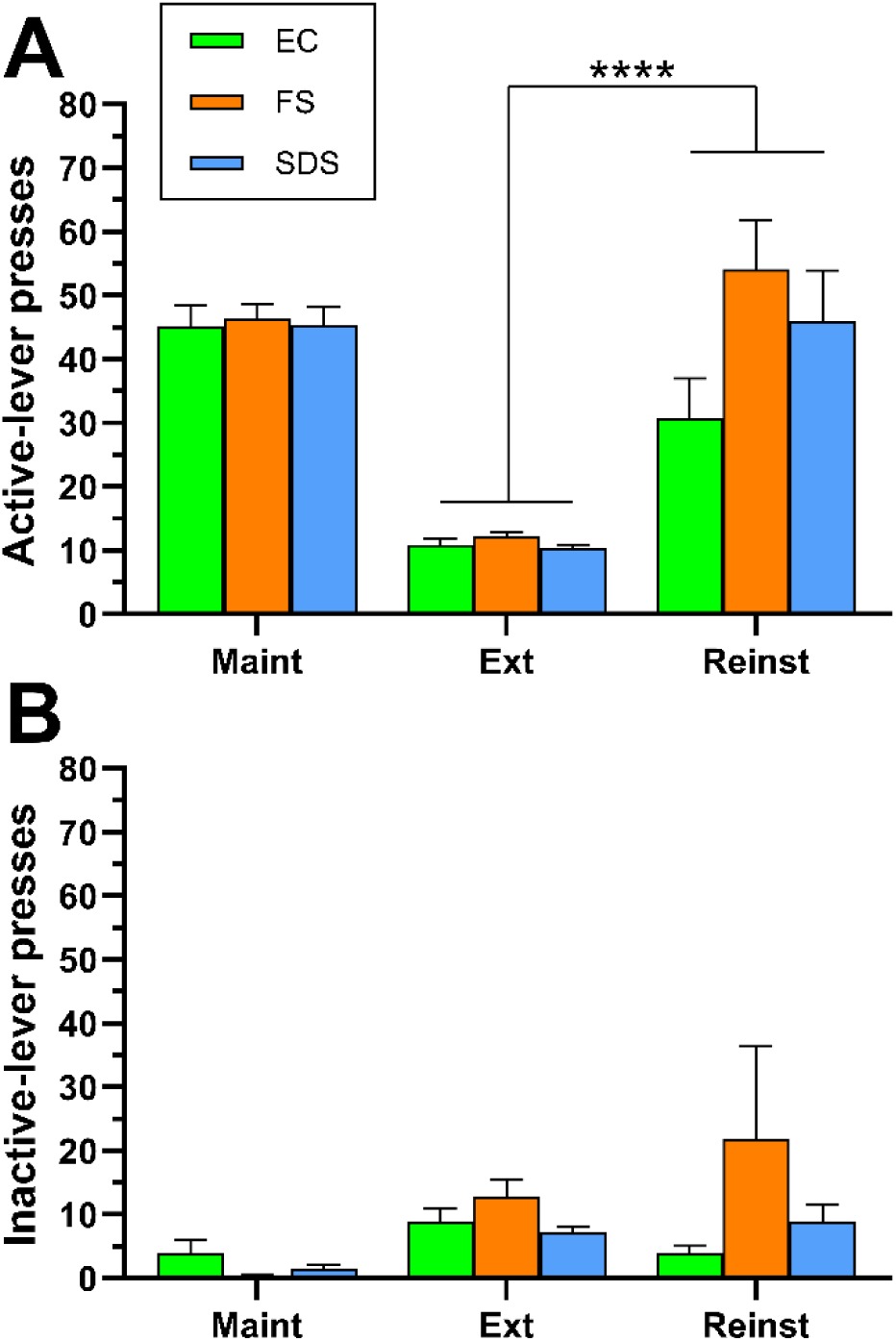
Reinstatement of cocaine-seeking behavior in SDS, FS, and EC rats. Shown are levels of responding on the **(A)** active lever and **(B)** inactive lever during the final 3 sessions of the maintenance phase of cocaine self-administration (Maint, sessions 8-10), the final 3 extinction sessions (Ext), and a reinstatement test session in which animals were re-exposed to a tactile cue that signaled impending social defeat stress (SDS, n=16, 8M/8F), footshock (FS, n=12, 6M/6F), or an empty cage control condition (EC, n=12, 6M/6F). Data are presented as mean ± SEM values. **** *p* < 0.0001, main effect of extinction vs. reinstatement (collapsed across groups).

### 3.5 Fos expression in VMH and PAG

We next analyzed Fos expression within the VMH and rostral/intermediate PAG (rPAG), two critical nodes within the medial hypothalamic defense system (**Fig. 5**). Because the majority of cocaine-seeking behavior occurs during the first 30 min of the reinstatement test session [102] (**Supplementary Fig. 3**), we elected to assess Fos expression patterns that would coincide with this period of maximal cocaine-seeking behavioral output. Activity-induced Fos protein expression typically peaks at 60-120 min post-stimulus [46,115,116], therefore we harvested brains at the immediate conclusion of the 2-h reinstatement test session, i.e. 90 min after the 30-min period of maximal cocaine seeking had occurred.

**Figure 5:**
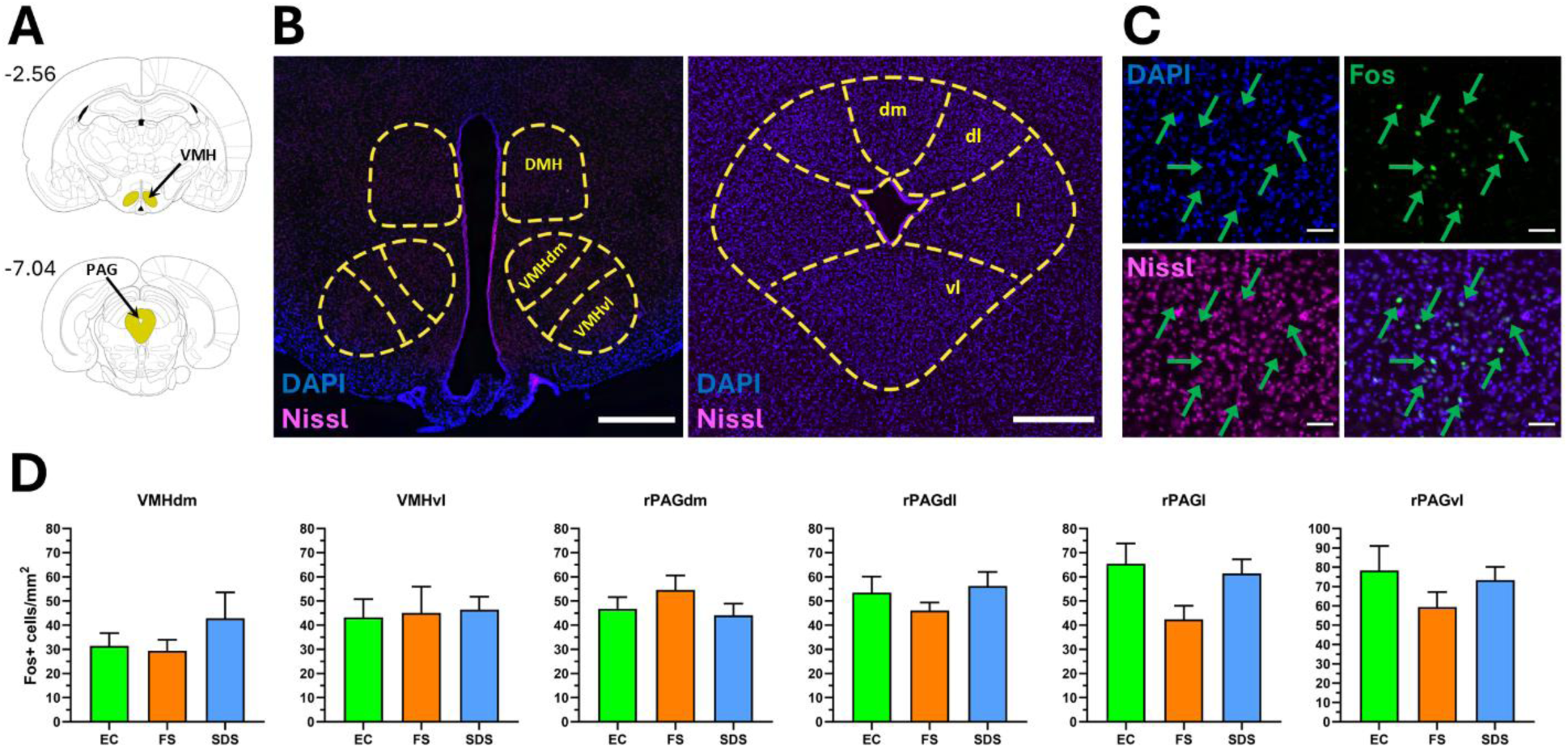
Fos expression in VMH and rPAG subregions during reinstated cocaine-seeking behavior does not differ between SDS, FS, and EC rats. Representative **(A)** rat brain atlas [114] images and **(B)** coronal section photomicrographs delineating boundaries of VMH and rPAG subregions. Numbers adjacent to atlas images in **A** indicate targeted A/P coordinates from Bregma. Mosaic photomicrographs in **B** were counterstained with DAPI (blue) and Nissl stain (magenta) and stitched together from a series of images initially acquired at 20x magnification. **(C)** Representative 40x images from rPAG containing DAPI (blue), Nissl stain (magenta), and Fos-positive cells (green). Green arrows point to examples of Fos-positive cells. **(D)** Quantification of mean (± SEM) Fos-positive cells/mm^2^ in VMH and rPAG subregions (n=11-16 rats/group). Scale bars represent **(B)** 500 μm or **(C)** 50 μm. DMH, dorsomedial hypothalamus. VMHdm and VMHvl, dorsomedial and ventrolateral aspects of ventromedial hypothalamus, respectively. dm/rPAGdm, dorsomedial rPAG. dl/rPAGdl, dorsolateral rPAG. l/rPAGl, lateral rPAG. vl/rPAGvl, ventrolateral rPAG.

Within each experimental group (EC, FS, SDS), no sex differences in Fos expression were detected for any VMH or rPAG subregion (*p* > 0.05, data not shown), therefore data from male and female subjects were combined. One-way ANOVAs for each of the VMH and PAG subregions found no significant differences in Fos expression between the three experimental groups (**Fig. 5D**), though there was a strong trend for the rPAGl which narrowly missed significance and appeared to be driven by greater levels of Fos in the EC and SDS groups as compared to the FS group (VMHdm, *F*_2,36_ = 0.78, *p* = 0.466; VMHvl, *F*_2,36_ = 0.05, *p* = 0.954; rPAGdm, *F*_2,37_ = 1.05, *p* = 0.361; rPAGdl, *F*_2,37_ = 0.86, *p* = 0.431; rPAGl, *F*_2,37_ = 3.20, *p* = 0.053; rPAGvl, *F*_2,37_ = 1.07, *p* = 0.354).

Although significant differences in mean Fos expression were not detected between the experimental groups, we considered the possibility that cocaine-seeking behavior in each group might be differentially correlated with neural activity in distinct VMH or rPAG subregions. We therefore performed a PCA separately for each experimental group to determine whether Fos expression in any VMH or rPAG subregion was associated with cocaine-seeking magnitude (i.e., number of active-lever presses during reinstatement). For the SDS group, we also included “active-defense” and “passive” behavioral scores as variables of interest in the PCA, as behaviors within these categories were previously found to correlate nearly or significantly with cocaine-seeking magnitude [102].

For the SDS group, PCA extracted three factors with an eigenvalue > 1. Cocaine-seeking magnitude loaded onto factor 1 and was accompanied by three other variables: rPAGl Fos expression, rPAGvl Fos expression, and “active-defense” score (*r* = 0.66 – 0.85). These variables did not cross-load onto the other two extracted factors, as factor 2 consisted of Fos expression in rPAGdm, rPAGdl, and VMHdm (*r* = 0.65 – 0.90), and factor 3 consisted of VMHvl Fos expression and “passive” score (*r* = 0.76 – 0.87). For the FS group, the Bartlett’s test of sphericity yielded a *p* value of *p* = 0.35, indicating that the data did not significantly differ from an identify matrix and was thus not suitable for PCA data reduction, i.e. no significant associations among the included variables could be extracted. For the EC group, PCA extracted two factors with an eigenvalue > 1. Cocaine-seeking magnitude loaded onto factor 1 and was accompanied by Fos expression in the rPAGdl, rPAGl, and rPAGvl (*r* = 0.77 – 0.88). These variables did not cross-load onto factor 2, which consisted of Fos expression in rPAGdm, VMHdm, and VMHvl (*r* = 0.62 – 0.82).

These unbiased PCA results indicated that at least for the SDS and EC groups, neuronal activity in one or more rPAG subregions, but not VMH subregions, may be a shared correlate of cocaine-seeking output. However, it was not possible from the PCA results to ascertain whether cocaine-seeking magnitude is more strongly associated with activation in any one rPAG subregion over another. To explore this question more deeply, subsequent two-variable correlation analyses were performed to directly assess associations between cocaine-seeking magnitude and Fos expression in each rPAG subregion. This analysis revealed that cocaine seeking in SDS and EC groups was associated with neural activity in distinct rPAG subregions. For the SDS group, reinstatement magnitude was significantly correlated with Fos expression only in the rPAGl (*r*_(14)_ = 0.61, *p* = 0.014), whereas reinstatement magnitude in the EC group was significantly correlated with Fos expression in the rPAGdl (*r*_(10)_ = 0.58, *p* = 0.047) and rPAGvl (*r*_(10)_ = 0.59, *p* = 0.044), but not with the rPAGl (**Table 1**). No rPAG subregion’s Fos expression was correlated with cocaine seeking in the FS rats, consistent with the null PCA results in this group.

**Table 1:** Correlation analyses of Fos expression in rPAG subdomains with reinstatement magnitude.

| Group | PAG subregion | r | p value |
| --- | --- | --- | --- |
| *SDS* | rPAGdm | -0.02 | 0.936 |
|  | rPAGdl | -0.28 | 0.299 |
|  | **rPAGl** | **0.60** | **0.014\*** |
|  | rPAGvl | 0.35 | 0.180 |
| *FS* | rPAGdm | 0.51 | 0.093 |
|  | rPAGdl | 0.27 | 0.393 |
|  | rPAGl | 0.34 | 0.286 |
|  | rPAGvl | 0.42 | 0.176 |
| *EC* | rPAGdm | -0.02 | 0.954 |
|  | **rPAGdl** | **0.58** | **0.047\*** |
|  | rPAGl | 0.43 | 0.160 |
|  | **rPAGvl** | **0.59** | **0.044\*** |

Significant correlations are in bold.
n=11-16 subjects per correlation analysis.
\* $p < 0.05$ .

### 3.6 Fos expression in afferent sources of input to rPAG

Given that our PCA and correlation analyses described above identified neural activity within several rPAG subregions as correlates of cocaine-seeking behavior, we wondered if activity in these rPAG subdomains might also be correlated to activity in other brain regions which have previously been implicated in reinstatement neurocircuitry and have known afferent connections to the rPAG. We opted to focus on five brain regions of interest: plPFC, ilPFC, LH/PfA, VTA, and CeA [60,85,117]. Within each experimental group (EC, FS, SDS), sex differences in Fos expression were not detected for any of the PAG-projecting brain regions examined (Bonferroni-corrected t-tests, *p* > 0.05, data not shown), therefore data from male and female subjects were combined within each group for subsequent analyses. One-way ANOVAs comparing EC, FS, and SDS groups detected significant differences in Fos expression for the VTA (*F*_2,37_ = 6.54, *p* = 0.004) and for both orexin-positive and orexin-negative cell-types in the LH/PfA (orexin-positive cells, *F*_2,37_ = 5.78, *p* = 0.007; orexin-negative cells, *F*_2,37_ = 15.35, *p* < 0.0001) (**Fig. 6**). For each of these analyses, *post hoc* Holm-Sidak’s tests revealed that Fos expression was higher in the FS group as compared to either the EC or SDS groups. No group differences were observed for any of the other brain regions analyzed (CeA, *F*_2,35_ = 1.62, *p* = 0.212; plPFC, *F*_2,36_ = 0.30, *p* = 0.742; ilPFC, *F*_2,36_ = 0.02, *p* = 0.980) (**Fig. 6**).

**Figure 6:**
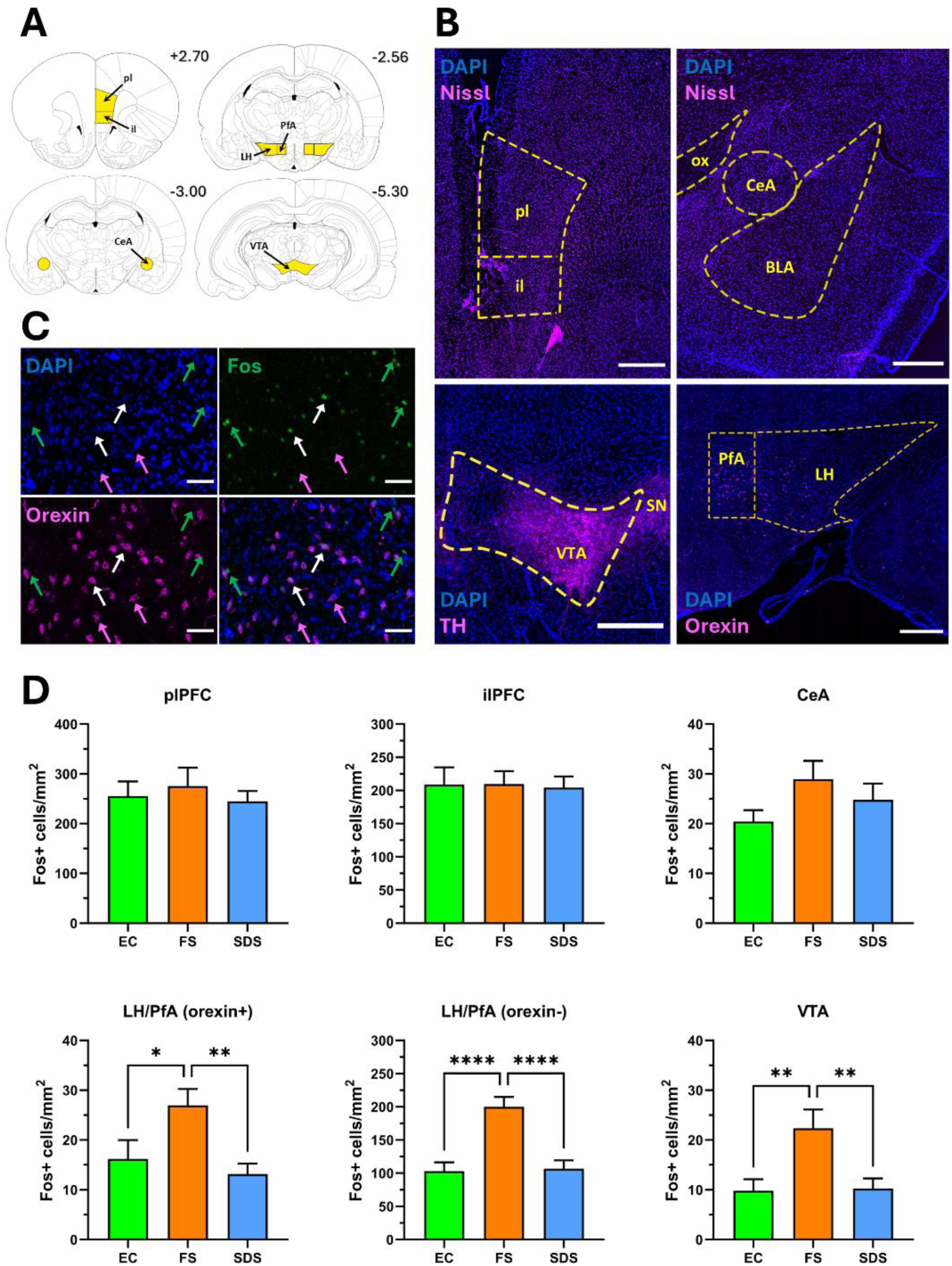
Levels of Fos expression within rPAG-projecting brain regions of interest during reinstatement tests in SDS, FS, and EC rats. Representative **(A)** rat brain atlas [114] images and **(B)** coronal section photomicrographs delineating boundaries for regions of interest (plPFC, ilPFC, CeA, LH/PfA, VTA). Numbers adjacent to atlas images in **A** indicate targeted A/P coordinates from Bregma. Photomicrographs in **B** were counterstained with DAPI (blue) and stitched together from a series of images initially acquired at 20x magnification. Anatomical borders in these sections were further revealed (magenta) via counterstaining with Nissl (pl/ilPFC, CeA), or via immunofluorescent detection of orexin (LH/PfA) or tyrosine hydroxylase (VTA). **(C)** Representative 40x images of LH/PfA containing DAPI (blue), orexin (magenta), and Fos-positive cells (green). Green arrows point to examples of Fos+/orexin-cells, magenta arrows point to examples of Fos-/orexin+ cells, and white arrows point to examples of Fos+/orexin+ cells. **(D)** Quantification of mean (± SEM) Fos-positive cells/mm^2^ in regions of interest (n=11-16 rats/group). Scale bars represent **(B)** 500 μm or **(C)** 50 μm. pl/plPFC, prelimbic PFC. il/ilPFC, infralimbic PFC. ox, optic tract. CeA, central amygdala. BLA, basolateral amygdala. LH, lateral hypothalamus. PfA, perifornical area. VTA, ventral tegmental area. SN, substantia nigra pars compacta. TH, tyrosine hydroxylase.

We next assessed whether levels of Fos expression in these PAG-projecting brain regions were differentially correlated with Fos expression in rPAG subregions of EC, FS, or SDS rats during their respective reinstatement tests (**Table 2**). In the SDS group, Fos expression in the rPAGl (the PAG subregion which we previously found was significantly correlated with cocaine-seeking magnitude, **Table 1**) was positively correlated with Fos expression in the plPFC (*r*_(14)_ = 0.59, *p* = 0.017) and in non-orexinergic cells of the LH/PfA (*r*_(14)_ = 0.59, *p* = 0.017). In EC rats, Fos expression in rPAG subdomains that were associated with cocaine seeking in this group (rPAGdl, rPAGvl) showed significant correlations with Fos expression in plPFC (rPAGdl and plPFC, *r*_(10)_ = 0.71, *p* = 0.010; rPAGvl and plPFC, *r*_(10)_ = 0.78, *p* = 0.003), ilPFC (rPAGvl and ilPFC, *r*_(10)_ = 0.72, *p* = 0.009), orexinergic cells in the LH/PfA (rPAGdl and LH/PfA orexin-positive cells, *r*_(10)_ = 0.68, *p* = 0.016), and CeA (rPAGvl and CeA, *r*_(10)_ = 0.58, *p* = 0.048). Fos expression in the VTA did not correlate with Fos expression in rPAG subregions that had previously been associated with cocaine-seeking behavior in either SDS or EC rats, and no significant correlations were detected in the FS group (**Table 2**).

**Table 2:**
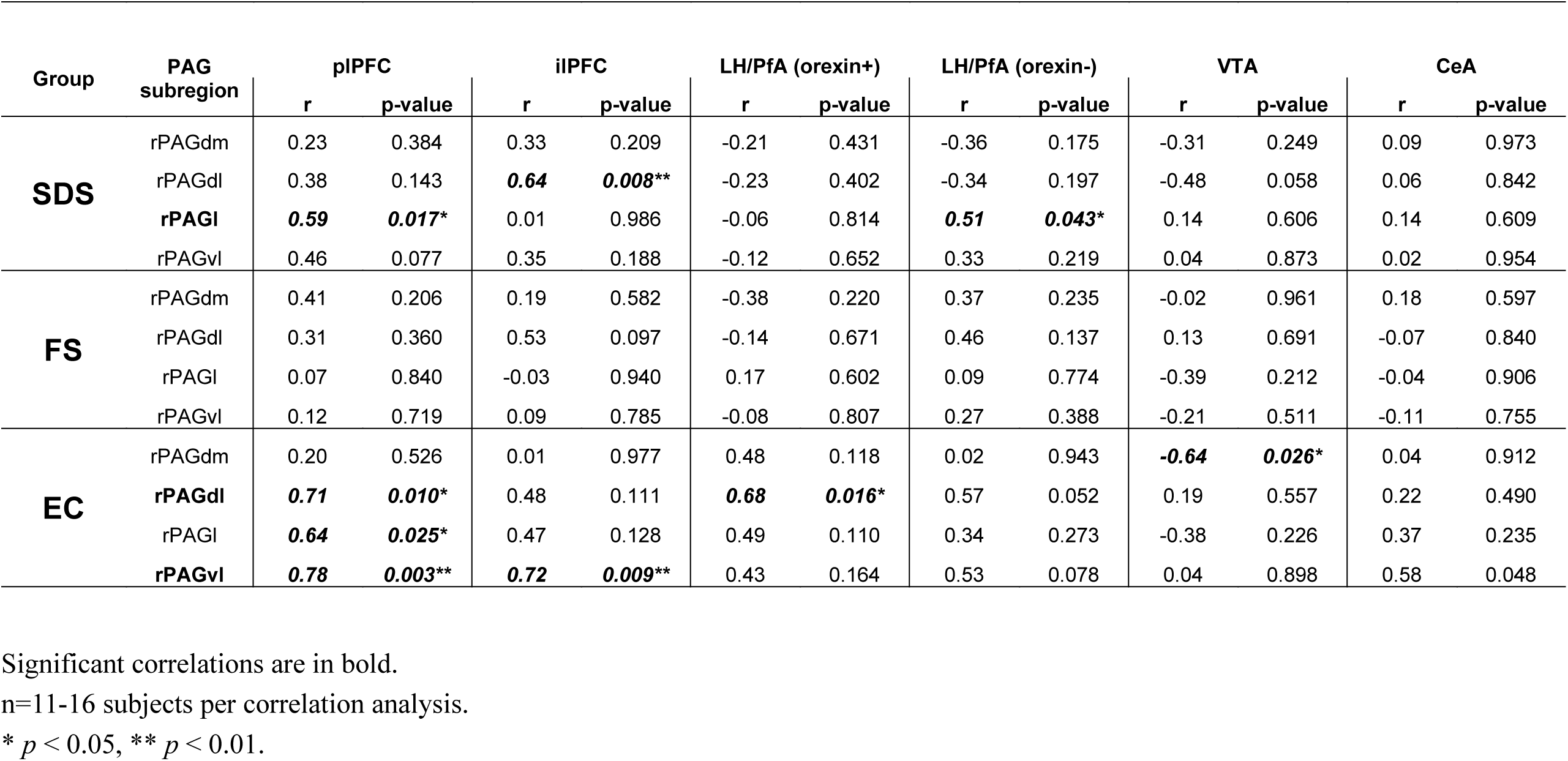
Correlations between Fos expression in PAG subregions and selected sources of afferent input.

### 3.7 Sex differences in behavioral and neuroactivational correlates of cocaine seeking in SDS rats

Finally, to examine potential sex differences in behavioral or neuroanatomical correlates of psychosocial stress-induced cocaine-seeking behavior, relationships between variables that were identified by the preceding PCA and Fos correlational analyses to be associated with cocaine-seeking behavior in the SDS group were reanalyzed separately in SDS males and SDS females (n= 8/sex) (**Fig. 7, Supplementary Table 1**). Correlation network analysis revealed a set of nodes in SDS males that showed positive interrelated correlations with cocaine-seeking behavior (**Fig. 7**). These included the behavioral variable extracted in the PCA (“active-defense” coping) as well as Fos expression in the rPAGl, plPFC, and non-orexinergic cells of the LH/PfA. No aspect of this correlation network was observed in SDS females, and no variables were found to correlate with cocaine seeking in the SDS females.

**Figure 7:**
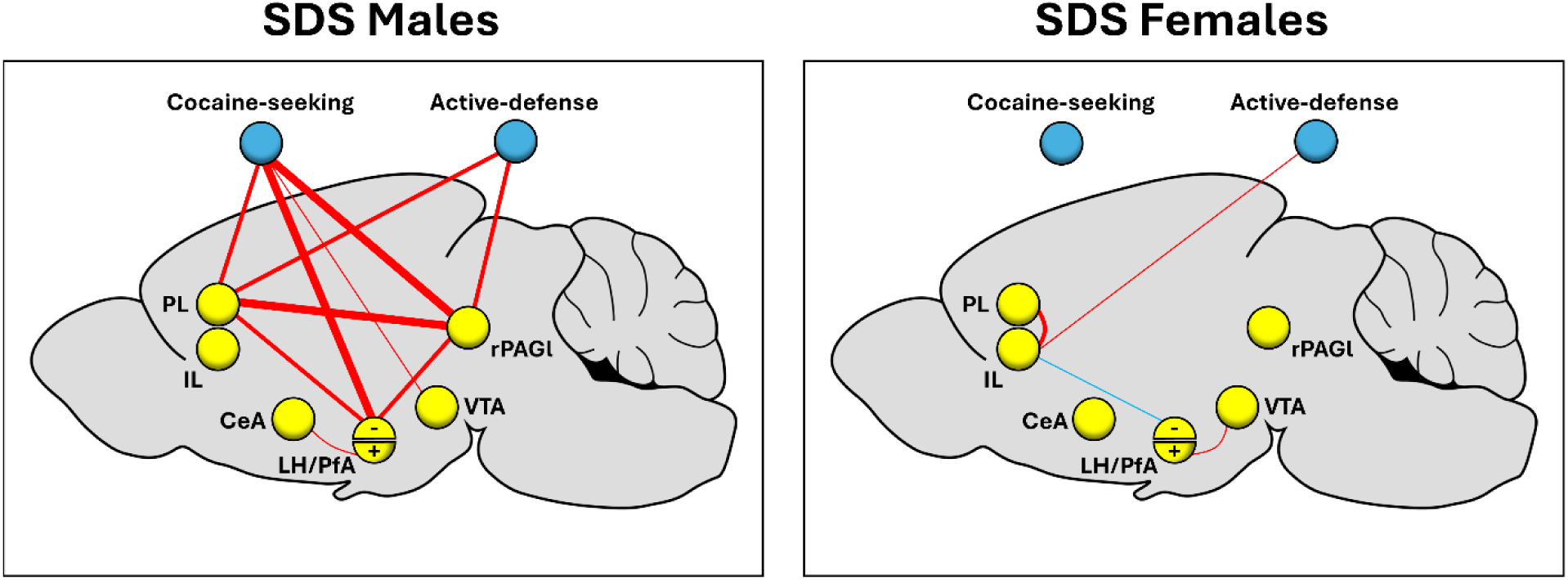
Schematic representation of correlation network analyses in SDS male (n=8) and SDS female (n=8) rats. Pearson’s correlations were used to assess potential associations between levels of reinstated cocaine-seeking behavior, “active-defense” scores derived from SDS episodes, and Fos expression in rPAGl and several of its afferent sources of input during cocaine-seeking behavior. Orexinergic and orexin-negative cells within the LH/PfA are designated by a “+” and “- “, respectively. Positive and negative Pearson’s *r* coefficient values are depicted as red and blue lines, respectively. Line weights (1, 3, 6 pt) represent Pearsons’s *r* coefficients as follows: 1 pt, *r =* ± 0.60 – 0.69; 3 pt, *r =* ± 0.70 – 0.79; 6 pt, *r =* ± 0.80 – 1.0. Nodes depict behavioral (blue) and neuroactivational (yellow) measures. n=7-8 for each two-variable correlation analysis.

## 4. DISCUSSION

In the present study, we replicated our previous finding that re-exposure to a discrete cue predictive of impending conspecific SDS reinstates cocaine seeking in male rats [102] and further demonstrated that this reinstatement response also occurs in females. In our initial characterization of this model, the SDS-predictive cue was a compound stimulus comprised of a tactile component (wire-mesh walls/floor) and an olfactory component (peppermint odor) [102]. However, we adjusted the parameters of the conditioned cue in the present study in two ways. First, we removed odor as a component of the stimulus because we have found it challenging in our current equipment configuration to limit its exposure to specific operant chambers without suffering some diffusion throughout the testing room or to other nearby operant chambers. Second, we altered the tactile cue such that it is now constructed of heavyweight durable polycarbonate rather than thin-gauge wire mesh, the former of which is less prone to damage, maneuvering, or manipulation by the resident-aggressor during protected-contact periods of SDS episodes. It is important to note that the redesigned SDS-predictive cue retained its capacity to engender cocaine seeking despite these parametric adjustments.

We also extended our earlier work by demonstrating that re-exposure to a cue predictive of impending uncontrollable footshock also elicits a robust cocaine-seeking response in rats that is similar in magnitude to levels of cocaine seeking produced by an SDS-predictive cue. Inclusion of this experimental group in the present and future studies will thus allow for meaningful comparisons of neuroactivational or other measures between subjects undergoing psychosocial vs. nonsocial stress-induced drug seeking responses. The reinstatement effect we observed here in response to a footshock-conditioned tactile cue is consistent with a previous study in which an auditory stimulus conditioned to footshock reinstate alcohol seeking in male rats [118], although others have reported that a footshock-conditioned auditory cue fails to induce reinstatement of heroin or cocaine seeking [17]. However, there are several important procedural differences that might account for these discrepancies. Most notably, we (present study) and others [118] have found that footshock-conditioned cues elicit reinstatement when footshock-cue conditioning takes place during or after the drug self-administration phase of the experiment, but not when conditioning takes place in drug-naïve animals prior to self-administration training [17]. Drug self-administration training/exposure may therefore have to precede stress conditioning in order for stress-conditioned cues to subsequently reinstate drug-seeking responses. This could be an important methodological consideration when the experimental objective is to engender stress-induced relapse-like operant behavior via the presentation of stress-conditioned stimuli.

The detection of a significant reinstatement effect in the empty cage (EC) control group was somewhat unexpected since the tactile cue presented to these animals during their reinstatement test session had not previously been associated with any overt stressor, and this same experimental manipulation induced a modest, albeit nonsignificant, level of cocaine seeking in our prior work [102]. As originally suggested in that study, we maintain that the cocaine-seeking response observed in this unstressed control group is most likely the result of an association between the tactile cue and cocaine availability that is not extinguished under the present testing conditions. Consistent with this view, only cocaine seeking induced by an SDS-predictive cue (and not an empty cage-predictive cue) was accompanied by a significant rise in serum corticosterone, indicative of a concurrent physiological stress-like state only in the SDS animals [102]. Although serum corticosterone was not analyzed in the present study, we would hypothesize that corticosterone levels would have similarly increased during reinstatement testing in the SDS and FS groups, but not in the EC group. It will be important in future studies to test this supposition in order to support or refute the notion that reinstatement in the SDS and FS groups, but not the EC group, is associated with induction of a stress-like state. An alternative possibility worth considering is that the tactile cue, functioning as a discriminative stimulus for cocaine availability, evokes a modest reinstatement effect in each of the three experimental groups, and this reinstatement effect is potentiated in the SDS and FS groups due to the cue’s secondary association with stressor exposure. This view is in line with previous evidence that stressor exposure enhances the reinstatement-inducing effects of drug-paired stimuli in animals [119–121] and cue-induced cocaine craving in cocaine-dependent human subjects [122,123]. In fact, some studies have suggested that stressors cannot elicit reinstatement if drug-conditioned cues are not present during reinstatement testing [16,124,125]. Future studies will be needed to determine whether the tactile cue’s association with cocaine availability in our experimental model contributes to the cocaine-seeking responses observed in the SDS- and FS-conditioned animals. Nevertheless, we offer that the EC group is a useful control for comparing neural activation patterns with SDS and FS groups since these animals exhibit cocaine-seeking responses (and accompanying neuroactivational patterns) that are unlikely to involve a stress-like state.

A major goal of this study was to test the hypothesis that neural activation within components of the medial hypothalamic defensive system would be selectively associated with psychosocial stress-induced cocaine-seeking responses in the SDS group as compared to the nonsocial-stress (FS) and EC groups, as these latter two cohorts exhibit cocaine-seeking behavior in the theoretical absence of psychosocial threat. Given the well-established role of the VMHvl in mediating defensive reactions to conspecific SDS [34,40,42,50–52,54], we were surprised to find no difference in Fos expression in this region between the experimental groups, nor was VMHvl activity associated with cocaine-seeking magnitude in the SDS group as determined by the PCA. Based on our results, we cannot presently attribute any potential contribution of the VMHvl to cocaine-seeking behavior, however it should be noted that Fos expression as a marker of neuronal activation may under some circumstances fail to identify a functional role for the VMH in defensive coping behaviors. For example, exposure to a context previously associated with predator threat does not increase Fos expression in the neighboring VMHdm, yet defensive responses to the predator-associated context are attenuated following chemogenetic inhibition of the VMHdm, suggesting that it is necessary for appropriate behavioral responsivity to a psychosocial stress-associated context even despite the lack of a detectable increase in Fos expression [45]. It is plausible therefore that the VMHvl may similarly play a functional role in cocaine seeking elicited by the SDS-predictive cue, but that such involvement was obfuscated by the use of c-Fos expression as a marker of neural activity in the present work. Additional studies will be necessary to further probe the role of the VMHvl in psychosocial stress-induced cocaine-seeking behavior before its potential involvement can be fully ruled out, perhaps using different immediate early gene markers of neuronal activation, *in vivo* measurement of neural activity, and/or by testing the causal role of the VMHvl in cocaine seeking via optogenetic, chemogenetic, or pharmacological manipulations.

Although significant group differences in mean Fos expression were also not detected within any of the rPAG subregions, several correlations between rPAG subregional neural activity and cocaine-seeking magnitude emerged from the data analysis, and the patterns of these correlations were distinct between the SDS, FS, and EC experimental groups. In particular, Fos expression within the rPAGl was positively and uniquely correlated with cocaine-seeking magnitude in the SDS group, i.e. greater levels of cocaine seeking elicited by impending conspecific social defeat coincided with greater neural activation in the rPAGl. To our knowledge, these are the first preclinical results identifying PAG activation as a correlate of drug-seeking behavior, although this finding is in agreement with a small number of neuroimaging studies in humans reporting that PAG activation may be associated with increased cocaine craving [99,100]. The potential involvement of the rPAGl in psychosocial stress-induced cocaine seeking would align with its known role in mediating aggressive/defensive responses to conspecific social threat [37,82,85,86,126] and appetitive actions like predatory hunting [85,127–133]. Moreover, the rPAGl is known to be interconnected with several brain regions critical for the reinstatement of drug-seeking behaviors including the plPFC, LH, VTA, CeA, and BNST [60,85,117,127,134], and is thus anatomically well-positioned to coordinate drug seeking in response to perceived psychosocial distress. Indeed, our network analysis revealed that rPAGl Fos expression was positively correlated with Fos expression in the plPFC and LH/PfA during reinstatement, suggesting that rPAGl activation may be functionally coupled with established components of stress-induced reinstatement neurocircuitry during bouts of cocaine seeking. Because the dorsal aspects of the PAG (including the rPAGl) orchestrate defensive and physiological responses to conspecific social threats, it is intriguing to consider that the rPAGl helps drive cocaine-seeking behavior in the SDS group as a proactive defensive response to the negative affective state produced by the SDS-predictive cue. This stress-induced cocaine-seeking defensive reaction may have been acquired through the animals’ prior experiences self-administering cocaine in the presence of this cue during self-administration sessions 11, 14, 17, and 20. However, this remains purely speculative and further research will be needed to adequately test this hypothesis. Nevertheless, support for this view comes from our finding that male rats’ tendency to emit active-defensive behavioral responses during conspecific SDS episodes was positively associated later with rPAGl activation during the reinstatement test. It is important to note that in both our prior and current experiments, measurements of active-defense coping scores during SDS typically preceded reinstatement test sessions by days-to-weeks as they were temporally separated by extinction sessions, suggesting that the association of active-defense coping with greater cocaine-seeking magnitude (Manvich et al 2016) and concurrent rPAGl activation (present results) might represent an enduring and stable trait. One possible mechanistic explanation could be that the innate excitability of the rPAGl differs between individual animals. If true, then one could hypothesize that any behavioral action which is dependent upon rPAGl activation might be exhibited to greater effect in rats with a more excitable rPAGl as compared to their counterparts. Such behaviors might include active-defensive reactions to conspecific social threat, psychosocial stress-induced cocaine seeking, or stress-induced potentiation of reinstatement elicited by cocaine-paired cues, among others. Further studies will be needed to ascertain whether rPAGl excitability varies across subjects (or between sexes) and if this might dually underlie individual differences in active-defense and/or cocaine-seeking behavioral responses.

While rPAGl activation was not correlated with cocaine seeking in the FS or EC groups, cocaine-seeking magnitude in the EC group was significantly correlated with Fos expression in the rPAGdl and rPAGvl. It is unclear why neural activity in these regions might be correlated with drug seeking in EC rats, especially in the absence of overt threat. However, it suggests that neural activity in the rPAG scales with drug-seeking behavior elicited by various stimuli, including discriminative stimuli predicting cocaine availability, and perhaps in a subregion-specific manner. The rostral/intermediate aspects of the dorsolateral and ventrolateral PAG are known to exhibit differential patterns of afferent and efferent connectivity [60,85], and thus their association with cocaine seeking in EC rats may reflect influences from different neural systems that contribute to varying and perhaps nonoverlapping aspects of the cocaine-seeking response via distinct ascending and/or descending output projections. Although the ventrolateral PAG is canonically activated during periods of passive coping responses to threat [37,82,86], we do not believe that reinstatement in EC animals was accompanied by a stress-like state because 1) PAGvl-mediated passive coping responses would likely have been associated with suppressed, rather than higher, levels of operant cocaine seeking, and 2) the cocaine-seeking response in EC rats occurs in the absence of increased circulating levels of corticosterone [102]. In contrast to the SDS and EC groups, no rPAG subregion’s Fos expression was significantly correlated with cocaine seeking in the FS group, despite the fact that these rats exhibited robust reinstatement. This may indicate that reinstatement elicited by footshock-conditioned cues is independent of PAG activity, although this question would be best addressed by directly manipulating PAG activation during the FS reinstatement condition.

Fos expression during cocaine seeking was significantly higher in the VTA and LH/PfA of FS rats as compared to either EC or SDS rats, consistent with increased Fos expression observed in these regions in response to footshock-conditioned cues or contexts [135–137] (but see [138] for negative results in VTA). It is unclear why the FS animals exhibited higher neural activation in these regions as compared to SDS or EC animals when there was no significant difference in the magnitude of cocaine seeking between the three groups. One possible explanation is that the FS-predictive cue may have been a more salient stimulus than the SDS-predictive or EC-predictive cues and therefore induced greater levels of neural activation that were more conducive to enhanced c-Fos expression, but which were well above a threshold necessary for contributing to drug-seeking behavioral output. Interestingly, while Fos expression in LH/PfA of SDS animals was lower than that of the FS group, neural activation of LH/PfA orexin-negative cells was strongly correlated with cocaine-seeking behavior in SDS male rats. This raises the fundamental question as to what is most important when interpreting data derived from Fos expression analyses – differences in overall mean Fos expression between groups, or group differences in the patterns of correlations between neural activity and behavioral measures? Numerous studies can be referenced to support either option and there is likely no clear answer. For our purposes, we interpret the strong correlation between Fos expression in LH/PfA non-orexinergic cells and cocaine-seeking magnitude in male SDS rats as indicative of a potentially novel and functional role for this neuronal subpopulation in psychosocial stress-induced cocaine seeking, seemingly in a sex-dependent manner, that warrants further investigation. Although most research in the LH and PfA has focused on orexinergic cell populations and their functional roles in drug seeking or attentional/arousal processes, there is prior evidence for recruitment of non-orexinergic cells in cocaine- and ethanol-seeking responses [139,140]. Moreover, several studies have demonstrated increased Fos expression in the LH and/or PfA in response to conspecific social defeat stress or associated cues, however the chemical phenotype of the activated cells was not confirmed [141–144]. We do not know the neurochemical identity of the Fos-positive non-orexinergic cells in the present study, but the LH and PfA are heterogeneous brain regions known to contain neurons releasing glutamate, GABA, and melanin-concentrating hormone in addition to orexin [145]. Additional studies will be needed to determine the chemical phenotype of Fos-positive neurons in LH/PfA whose activation is correlated with psychosocial stress-induced cocaine seeking.

The PAG is interconnected with several brain regions implicated in stress-induced drug seeking processes. In the present study, we selected five of these brain regions as candidates for further Fos expression analyses because we hypothesized that they may influence PAG activity during cocaine seeking: plPFC, ilPFC, LH/PfA, VTA, and CeA [85]. Although there were no group differences in plPFC or ilPFC Fos expression, we found that Fos expression in the plPFC was positively and significantly correlated with neural activity in PAG subregions whose Fos expression was associated with cocaine-seeking magnitude in SDS and EC rats. More specifically, plPFC Fos expression correlated with activity in the rPAGl of SDS rats and with the rPAGdl and rPAGvl of EC rats. It was also noteworthy that plPFC activity trended towards a significant correlation with the rPAGvl in SDS rats (p = 0.077) and significantly correlated with the rPAGl in EC rats. Taken together, these results suggest that there may be coordinated communication between the plPFC and several PAG subregions during cocaine seeking evoked by diverse stimuli. The plPFC, a component of the medial PFC (mPFC), plays an important role in maintaining behavioral inhibition and cognitive flexibility that are critical for the expression of situation-appropriate adaptive responses to diverse environmental stimuli. Repeated social defeat stress, which under some conditions confers enhanced motivation for cocaine [146–150], reduces functional connectivity between the mPFC and dPAG and induces a social avoidance phenotype that can be recapitulated by disinhibition of dPAG glutamatergic neurons via chemogenetic inhibition of mPFC input to dorsal and lateral aspects of the PAG [151]. PAG activation has also previously been linked to perceived social rejection and social distress in humans [152], and recent human neuroimaging studies in cocaine-dependent subjects have reported that cocaine use/craving is associated with PAG hyperactivation, sometimes resulting from apparent deficits in medial prefrontal cortical-dependent top-down inhibition [99,100]. Extrapolating these findings to our present results, the observed positive correlations between Fos expression in PAG subregions and in plPFC during psychosocial stress-induced cocaine-seeking behavior can be interpreted in at least two ways. First, it is possible that the positive correlation represents bottom-up excitatory signaling from PAG to plPFC, the hyperactivation of which may contribute to the emergence of cocaine-seeking behavior [99] perhaps in part through the plPFC’s canonical role in reinstated drug-seeking responses [20,153–155]. Alternatively, it is possible that the positive correlation results from a loss of top-down inhibitory control by the plPFC over the PAG, resulting in a disinhibited PAG that in turn contributes to the expression of cocaine-seeking behavior. In this latter scenario, the positive correlation may arise because increased plPFC activity during a state of enhanced PAG activation represents an attempt by the plPFC to dampen PAG activity, a pathway that might be rendered ineffective through a loss of functional connectivity within the mPFC→PAG projection as has been demonstrated in cocaine-dependent subjects [99,100] and in rodents following repeated social defeat stress [151]. Future work aimed at delineating the directionality of the positive correlation between plPFC and PAG neural activity during various modalities of reinstated cocaine-seeking behavior will be needed to evaluate these possibilities.

Fos expression in the LH/PfA was also significantly and positively correlated with Fos expression in distinct rPAG subregions of SDS and EC rats, and in a cell-type specific manner. In EC rats, Fos expression in LH/PfA orexinergic cells was positively correlated with Fos expression in the rPAGdl, a PAG subregion whose activation scaled with cocaine-seeking magnitude in this experimental group. It was also noteworthy that Fos expression in non-orexinergic LH/PfA cells very narrowly missed significance as a correlate of rPAGdl activity (p=0.052). The LH/PfA is reciprocally, albeit weakly, interconnected with the PAGdl [156–158], though the functional significance of these pathways remains largely unknown. However, there is a well-established role for LH/PfA orexin signaling in cue- and context-induced reinstatement of cocaine seeking, as exposure to cocaine-conditioned cues or contexts activates orexinergic neurons in the LH and PfA [159–161], and cocaine seeking in response to these stimuli is attenuated by systemic or intra-VTA orexin receptor antagonism [162–167]. In general, there is convergent evidence that the orexinergic projection arising from LH (and PfA in some studies) and terminating in the VTA is a critical substrate for cue-induced reinstatement of cocaine seeking [168,169], thus its potential involvement in reinstatement elicited by the cocaine-associated tactile cue in the EC rats should not be wholly unexpected. Additionally, activation of LH/PfA non-orexinergic neurons specifically projecting to VTA has also been reported during cue-induced reinstatement of cocaine seeking [139]. However, whether activation of rPAGdl (or any PAG subregion) is involved in cue- or context-induced reinstatement, and whether its contributions to this reinstatement effect might arise from either its excitatory input to LH/PfA orexinergic neurons or as a functional efferent output of LH/PfA neurons, has not been explored to our knowledge. Our finding that rPAGdl activation is positively correlated with cocaine seeking in EC rats, and that its activity scales with orexinergic (and nearly with non-orexinergic) cells within the LH/PfA, points to a potential role for communication between these regions as a contributor to drug-seeking responses elicited by cocaine-associated cues.

In SDS animals, rPAGl activation correlated significantly with Fos expression in non-orexinergic cells of the LH/PfA during cocaine seeking. No other correlations were detected in SDS animals between any rPAG subregion and either orexinergic or non-orexinergic cells in LH/PfA. As mentioned above, activation of VTA-projecting LH/PfA orexin-negative cells has been observed previously during cue-induced cocaine seeking [139], but we are unaware of prior studies examining their activation during stress-induced reinstatement. We did not determine the connectivity patterns of the Fos-positive/orexin-negative LH/PfA cells, thus we do not know if the Fos-positive LH/PfA cells that correlated with psychosocial stress-induced cocaine seeking here represent a VTA-projecting subpopulation, nor do we presently know the neurochemical phenotype of these cells. Such questions will need to be addressed by future experiments. The rPAGl and its anatomical connections with the LH have received attention in recent years as a potentially important contributor to other motivated appetitive behaviors in rodents, best characterized by its involvement in foraging/predatory behavior [60,85,132]. The rPAGl receives a dense projection from the medial and lateral hypothalamic regions that contain orexinergic neurons, including the LH, PfA, and dorsomedial hypothalamus [117,157,170]. The lateral PAG exhibits increased neural activity during predatory hunting in rodents [127,171], and optogenetic activation of a GABAergic projection from LH to lateral PAG elicits predation in mice [171]. These findings indicate that a LH→PAGl projection mediates prey seeking, however the connection between rPAGl and LH/PfA is bidirectional [127,172] and rPAGl lesions have been found to reduce hunting-induced Fos expression in orexinergic cells of the LH/PfA, indicative of a possible rPAGl→LH/PfA pathway that is also engaged during hunting in rodents [127]. Thus, while the precise directionality of communication between LH/PfA and rPAGl during predation is still not fully resolved, these and other studies have implicated the rPAGl as an important contributor to hunting, foraging, and reward approach in rodents [60,85]. Moreover, there is accruing evidence that these appetitive motivated behaviors may be driven at least in part by putatively non-orexinergic LH/PfA input to rPAGl [171]. When considered together with evidence that the rPAGl is also involved in defensive responses to social threat [37,82,85,86,126] and its activation was correlated with reinstatement magnitude in our current study, it seems plausible to hypothesize that psychosocial stress-induced cocaine seeking might be mediated by rPAGl activation that is either downstream or upstream of non-orexinergic LH/PfA activation.

Up to this point, male and female subjects were collapsed together in all behavioral and neuroactivational analyses, however we were curious as to whether any of the observed correlations between endpoints may have been more robustly driven by one sex over another. When we performed a correlation network analysis incorporating cocaine-seeking magnitude, active-defense coping scores during SDS episodes, and neural activity during cocaine seeking separately in SDS males and SDS females, a rather striking sex difference emerged. In males, we found a highly-interconnected network that positively associated levels of neural activity between rPAGl, plPFC, and orexin-negative LH/PfA cells. Moreover, Fos expression in each of these nodes also correlated with cocaine-seeking magnitude and active-defense during SDS. For most of these correlations, Pearson’s *r* coefficients were moderate-to-strong. By contrast, no aspect of this correlation network appeared in SDS female rats. In fact, other than a positive correlation between neural activity in plPFC and ilPFC, all remaining correlations examined in females were nonsignificant. It thus appears that associations between Fos expression in rPAGl, plPFC, and LH/PfA with cocaine seeking or active-coping selection in SDS subjects is largely driven by the males in the cohort. We cannot yet fully explain this sex difference in the network correlation analysis, although several possible explanations can likely be ruled out. For example, we did not find evidence for sex differences in cocaine self-administration or cocaine-seeking magnitude in any of our experimental groups including the SDS rats, thus it seems unlikely that these were contributing factors, although we note that our sample sizes were not necessarily powered to detect sex differences in these measures. The lack of sex differences in putative stress-induced cocaine seeking in the present study is in line with some, but not all, prior investigations, as only a small number of studies have explored sex differences in stress-induced reinstatement of drug seeking and have produced mixed results [16,20,173]. For example, some have reported that female rats exhibit greater stress-induced reinstatement of cocaine seeking than males [174–176], while others have found no obvious sex differences [177,178]. Additionally, stress-induced potentiation of cocaine-primed reinstatement via restraint stress or systemic administration of corticosterone is more or less equally effective in both sexes, although it was noted that footshock potentiates cocaine-primed reinstatement only in males [179]. By contrast, the combination of cocaine-associated cues and yohimbine produced higher levels of cocaine-seeking in females than males [177]. Given these equivocal results and a lack of sex differences in stress-induced cocaine-seeking magnitude in the present study, we do not believe that differential levels of cocaine seeking in response to the SDS-predictive cue between males and females (or differential sensitivity to stress-induced reinstatement of cocaine seeking in general) contributed to the male-driven correlation metrics. One could also speculate that the sex-dependent correlational patterns resulted from a reliance on different neural systems in males vs. females for coordinating action responses to impending conspecific social aggression, however most studies have indicated that the functions of the VMH, PMD, and PAG in producing defensive responses to psychosocial threats are equivalent between sexes across species [32,34,42,45,88,180–189]. Moreover, the role for rPAGl in predatory hunting (i.e., appetitive behavior) has also been replicated in female rodents [133].

Although the neural circuitry necessary to respond to psychosocial stressors is similar between males and females, we cannot be certain that the qualitative experience of conspecific same-sex aggression was equivalent between males and females in our study. Males are more aggressive than females during conspecific social interactions in many species [190], and in rats, attack behaviors emitted by females are generally less forceful or aggressive than those emitted by males [189]. We did not quantify resident-aggressor behaviors and thus cannot determine whether SDS episodes might have differed qualitatively between males and females, however we anecdotally observed that male-male interactions were more typically aggressive than female-female interactions, and the impacts of social defeat stress on various measures are known to sometimes differ between male and female rodents [150,191–194]. We therefore cannot rule out the possibility that the selective association of rPAGl, plPFC, and LH/PfA activation with active-defense coping and cocaine seeking in males is the result of more aggressive, physically-engaging, and/or salient SDS encounters as compared to their female counterparts. While male rats typically do not aggress towards unfamiliar females and instead display copulatory behaviors, female-directed attacks can be elicited in male rodents via electrical, optogenetic, or chemogenetic stimulation of the VMHvl [50,189,195,196]. This may provide a means for examining whether enduring more aggressive/salient conspecific aggression would produce neuroactivational correlates of cocaine seeking and/or active-defense behaviors in female subjects that are similar to what was observed in males.

This study has several limitations that should be mentioned. First, not all components of the medial hypothalamic defense system were examined in our Fos analyses. We selected the VMH and PAG as our primary seed regions of interest on the basis that 1) their activation is necessary for behavioral responsivity to conspecific social aggression in rodents [34,37,40,40,41,63], 2) they are easily identifiable using common immunohistochemical methods coupled with fluorescence microscopy, and 3) their capacity to engender fear-like responses is conserved in nonhuman primates [39,93] and humans [95–97]. However, we acknowledge that there are several additional hypothalamic nuclei known to be involved in the manifestation of defensive responses to psychosocial threats, including the dorsal and ventral premammillary nuclei, anterior hypothalamus, and medial preoptic area [38,40]. Second, our Fos expression analyses cannot inform whether the rPAG is causally involved in cocaine-seeking behavior since they are correlational in nature. Additional studies employing pharmacological, chemogenetic, optogenetic, or neuronal ensemble-targeted approaches will be required to determine whether activation in any rPAG subregion is necessary and/or sufficient for the expression of cocaine-seeking behavior. Third, all of our experimental subjects had a history of cocaine self-administration prior to reinstatement testing and Fos expression analysis, and we are therefore unable to ascertain whether this prior cocaine exposure might have affected Fos expression patterns in response to the SDS-, FS-, or EC-predictive cues. To our knowledge, no studies have examined whether prior psychostimulant exposure alters SDS-induced neuroactivational responses, however it is plausible given that prior exposure to SDS alters psychostimulant-induced patterns of neural activity in PAG and other brain regions in rodents [66,149]. Finally, we acknowledge that some correlational analyses may have been slightly underpowered. It is worth noting that neither of the correlation network analyses in SDS males or SDS females detected a significant correlation between active-defense behavior and cocaine seeking, although this may be due to insufficient statistical power as our previous study reporting a correlation between these variables was comprised of a larger sample size of male rats [102]. It will be important in future studies to corroborate the associations between cocaine-seeking magnitude, active-defensive coping, and neuroactivational patterns identified here in larger sample sizes of both male and female subjects.

## 5. CONCLUSIONS

In conclusion, we report that previously-extinguished cocaine-seeking behavior can be robustly reinstated by re-exposure to a discrete cue that is predictive of impending conspecific social aggression or footshock in male and female rats. Analysis of Fos expression patterns suggests a previously-unidentified potential role for the rPAG in emergent cocaine-seeking behavior, and for the rPAGl specifically in psychosocial stress-induced cocaine seeking in male rats. Finally, we show that rPAGl activation is positively correlated with cocaine-seeking magnitude in SDS-exposed rats, predilection to display active-defensive responses to conspecific social threat, and neural activity in plPFC and non-orexinergic cells of the LH/PfA, and that these associations are largely driven by males. Taken together with recent neuroimaging findings in human subjects, our results lend further support to the potential functional involvement of PAG activation in stress-induced drug-seeking behavior, potentially in a sex-dependent and subregion-specific manner.

## ACKNOWLEDGEMENTS

The authors would like to thank the Rowan-Virtua SOM Histopathology Core for the generous use of their tissue sectioning equipment.

## DATA STATEMENT

The datasets generated and analyzed in the current study are available from the corresponding author upon reasonable request.

