## Supplementary Material for "Fos expression in the periaqueductal gray, but not the ventromedial hypothalamus, is correlated with psychosocial stress-induced cocaine-seeking behavior in rats"

### Supplementary Methods and Materials

*Operational definitions of behaviors observed and scored during social defeat stress episodes*

Boxing – The subject stands upright on hind legs with forepaws outstretched and contacts opponents upper-torso, head, or forepaws.

Push – The subject purposefully places forepaws on any aspect of the opponent and initiates directed movement towards the opponent. Incidental contact between subject and opponent and opponent-initiated contacts are not considered a “push” by the subject.

Dominant Posture – The subject is situated on top of an opponent that has assumed the submissive supine posture. The forepaws typically rest somewhere along the torso or on the extended forepaws of the opponent. The hind legs may or may not also rest on the opponent.

Escape/Flight – The subject exhibits deliberate and high-velocity locomotion in no particular direction with an obvious intention of avoiding contact with the opponent. Longer durations (> 1 sec) of “escape” are typically expressed as circular thigmotaxis with occasional movements through the center of the cage. Attempts to avoid the opponent by means of jumping (whether in an open area or along the walls of the cage) are also scored as escape/flight behavior.

Upright Defense – The subject stands upright or slightly hunched on hind legs but otherwise remains still in a “ready-to-act” posture. Forepaws may be close to the body or outstretched. Subjects often track the movement of the opponent and may make small adjustments to their position seemingly to keep the opponent localized in front of them.

Freezing – The subject, in a standing position with all four paws contacting the ground, ceases all movement with the exception of breathing, slight reactions to contact by the opponent, and occasional tail flicks/rattles. Eyes may sometimes appear half-closed.

Supine Submissive Posture – The subject lies prone on their back, motionless, with paws extended upwards. The posture is most often observed when the opponent has “pinned” the subject and is standing in a dominant posture over the subject.

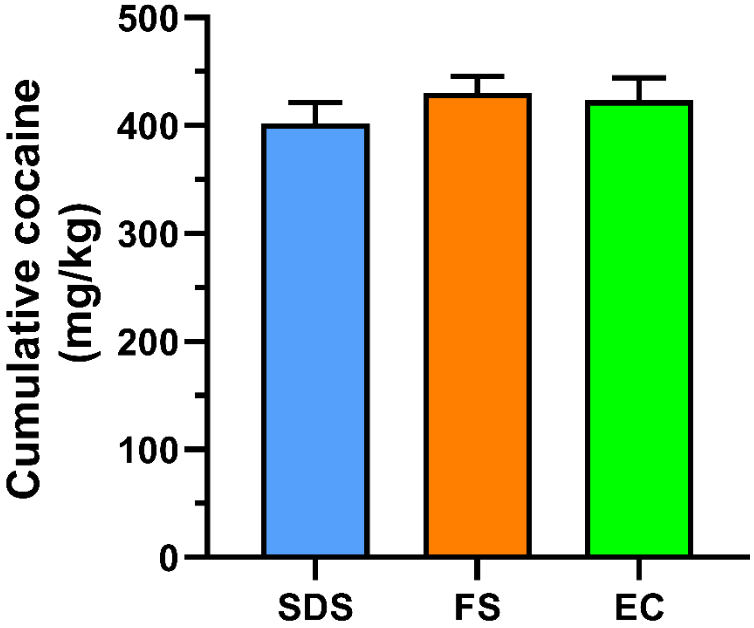


**Supplementary Figure 1. Cumulative cocaine intake resulting from IV cocaine self-administration does not differ between SDS, FS, and EC rats.** Rats self-administered 0.5 mg/kg/inf IV cocaine for 20 sessions under a FR1 schedule of reinforcement. Shown is mean ± SEM cumulative cocaine intake summed over the 20 cocaine self-administration sessions in the SDS (n=16, 8M/8F), FS (n=12, 6M/6F) and EC (n=12, 6M/6F) experimental groups.

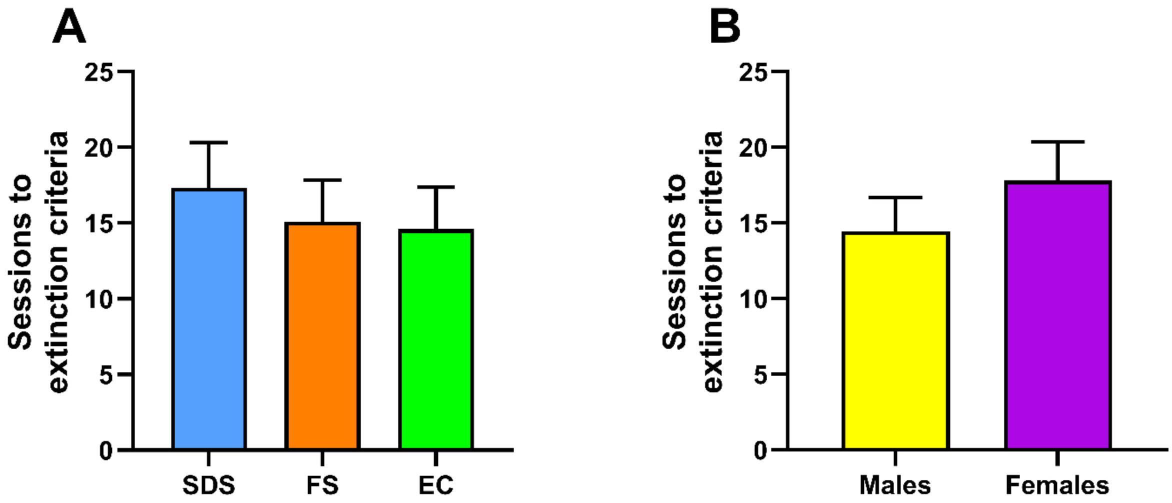


**Supplementary Figure 2. No group or sex differences in extinction training.** Following cocaine self-administration, responding was extinguished in daily 2-hr sessions until extinction criteria were satisfied (< 15 active-lever responses in 3/4 consecutive sessions). **(A)** Number of sessions required to achieve extinction criteria in SDS (n=16, 8M/8F), FS (n=12, 6M/6F) and EC (n=12, 6M/6F) rats. **(B)** Number of sessions required to achieve extinction criteria in male (n=20) and female (n=20) rats, collapsed across experimental groups. Data are presented as mean ± SEM values.

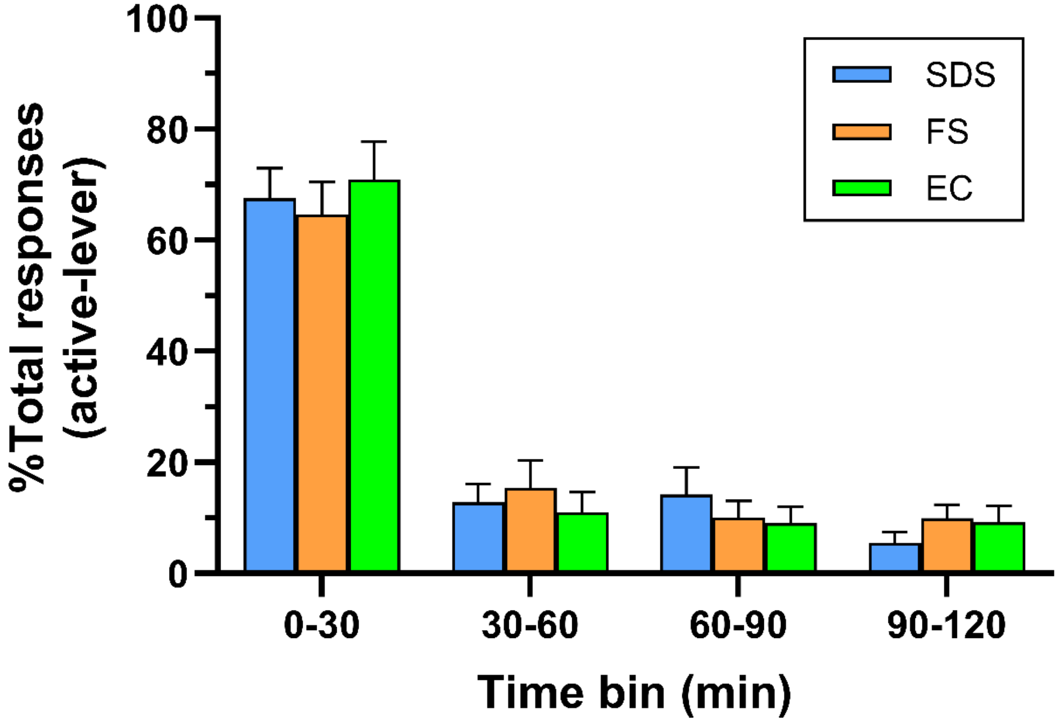


**Supplementary Figure 3. Distribution of cocaine-seeking behavior across the reinstatement test.** Shown are the percentages of active-lever responses across each 2-hr reinstatement test in 30-min bins in SDS (n=16, 8M/8F), FS (n=12, 6M/6F) and EC (n=12, 6M/6F) rats. For each group, the summation of percentages across bins totals 100%. Data are presented as mean ± SEM values.

***Supplementary Table 1: Correlation matrix in SDS male and SDS female rats***

| | Group | cocaine seeking | | "active-defense" | | rPAGl | | plPFC | | ilPFC | | LH/PfA (orexin+) | | LH/PfA (orexin-) | | VTA | | CeA | |
|---|---|---|---|---|---|---|---|---|---|---|---|---|---|---|---|---|---|---|---|
| | | r | p-value | r | p-value | r | p-value | r | p-value | r | p-value | r | p-value | r | p-value | r | p-value | r | p-value |
| **SDS Males** | cocaine seeking | --- | --- | 0.45 | 0.259 | ***0.81*** | ***0.015**** | ***0.76*** | ***0.029**** | 0.23 | 0.592 | 0.01 | 0.983 | ***0.84*** | ***0.009***** | 0.67 | 0.067 | 0.29 | 0.491 |
| | "active-defense" | 0.45 | 0.259 | --- | --- | ***0.76*** | ***0.030**** | ***0.74*** | ***0.035**** | 0.49 | 0.218 | 0.06 | 0.887 | 0.25 | 0.554 | 0.12 | 0.786 | 0.39 | 0.344 |
| | rPAGl | ***0.81*** | ***0.014**** | ***0.76*** | ***0.03**** | --- | --- | ***0.93*** | ***< 0.001****** | ***0.33*** | ***0.042**** | 0.17 | 0.695 | ***0.74*** | ***0.038**** | 0.38 | 0.351 | 0.35 | 0.402 |
| | plPFC | ***0.76*** | ***0.029**** | ***0.74*** | ***0.035**** | ***0.93*** | ***< 0.001****** | --- | --- | 0.55 | 0.155 | 0.09 | 0.832 | ***0.76*** | ***0.030**** | 0.37 | 0.363 | 0.34 | 0.412 |
| | ilPFC | 0.23 | 0.592 | 0.49 | 0.218 | 0.33 | 0.420 | 0.55 | 0.155 | --- | --- | -0.47 | 0.235 | 0.25 | 0.551 | -0.23 | 0.582 | 0.18 | 0.662 |
| | LH/PfA (orexin+) | 0.01 | 0.983 | 0.06 | 0.887 | 0.17 | 0.695 | 0.09 | 0.832 | -0.47 | 0.235 | --- | --- | 0.29 | 0.486 | -0.02 | 0.960 | 0.64 | 0.088 |
| | LH/PfA (orexin-) | ***0.84*** | ***0.009***** | 0.25 | 0.554 | ***0.74*** | ***0.038**** | ***0.76*** | ***0.030**** | 0.25 | 0.551 | 0.29 | 0.486 | --- | --- | 0.40 | 0.322 | 0.50 | 0.209 |
| | VTA | 0.67 | 0.067 | 0.12 | 0.786 | 0.38 | 0.351 | 0.37 | 0.363 | -0.23 | 0.582 | -0.02 | 0.960 | 0.40 | 0.322 | --- | --- | -0.24 | 0.570 |
| | CeA | 0.29 | 0.491 | 0.39 | 0.344 | 0.35 | 0.402 | 0.34 | 0.412 | 0.18 | 0.662 | 0.64 | 0.088 | 0.50 | 0.209 | -0.24 | 0.570 | --- | --- |
| **SDS Females** | cocaine seeking | --- | --- | 0.04 | 0.918 | 0.09 | 0.829 | 0.55 | 0.157 | 0.17 | 0.692 | 0.06 | 0.882 | 0.06 | 0.887 | 0.41 | 0.319 | 0.06 | 0.895 |
| | "active-defense" | 0.04 | 0.918 | --- | --- | -0.38 | 0.354 | 0.59 | 0.122 | 0.68 | 0.062 | -0.06 | 0.893 | -0.34 | 0.405 | -0.58 | 0.129 | 0.12 | 0.797 |
| | rPAGl | 0.09 | 0.829 | -0.38 | 0.354 | --- | --- | 0.24 | 0.564 | -0.24 | 0.567 | -0.26 | 0.535 | 0.43 | 0.286 | -0.10 | 0.806 | 0.08 | 0.869 |
| | plPFC | 0.55 | 0.157 | 0.59 | 0.122 | 0.24 | 0.564 | --- | --- | ***0.74*** | ***0.035**** | -0.22 | 0.601 | -0.32 | 0.441 | -0.39 | 0.336 | 0.46 | 0.304 |
| | ilPFC | 0.17 | 0.692 | 0.68 | 0.062 | -0.24 | 0.567 | ***0.74*** | ***0.035**** | --- | --- | -0.03 | 0.949 | -0.62 | 0.102 | -0.44 | 0.272 | 0.52 | 0.233 |
| | LH/PfA (orexin+) | 0.06 | 0.882 | -0.06 | 0.893 | -0.26 | 0.535 | -0.22 | 0.601 | -0.03 | 0.949 | --- | --- | 0.54 | 0.169 | 0.69 | 0.061 | 0.29 | 0.532 |
| | LH/PfA (orexin-) | 0.06 | 0.887 | -0.34 | 0.405 | 0.43 | 0.286 | -0.32 | 0.441 | -0.62 | 0.102 | 0.54 | 0.169 | --- | --- | 0.52 | 0.187 | -0.19 | 0.684 |
| | VTA | 0.41 | 0.319 | -0.58 | 0.129 | -0.10 | 0.806 | -0.39 | 0.336 | -0.44 | 0.272 | 0.69 | 0.061 | 0.52 | 0.187 | --- | --- | -0.10 | 0.830 |
| | CeA | 0.06 | 0.895 | 0.12 | 0.797 | 0.08 | 0.869 | 0.46 | 0.304 | 0.52 | 0.233 | 0.29 | 0.532 | -0.19 | 0.684 | -0.10 | 0.830 | --- | --- |

Significant correlations are in bold.
n=7-8 subjects per correlation analysis.
* $p < 0.05$, ** $p < 0.01$, *** $p < 0.0001$.
